# DeepResBat: deep residual batch harmonization accounting for covariate distribution differences

**DOI:** 10.1101/2024.01.18.574145

**Authors:** Lijun An, Chen Zhang, Naren Wulan, Shaoshi Zhang, Pansheng Chen, Fang Ji, Kwun Kei Ng, Christopher Chen, Juan Helen Zhou, B.T. Thomas Yeo, Alzheimer’s Disease Neuroimaging Initiative, Australian Imaging Biomarkers and Lifestyle Study of Aging

## Abstract

Pooling MRI data from multiple datasets requires harmonization to reduce undesired inter-site variabilities, while preserving effects of biological variables (or covariates). The popular harmonization approach ComBat uses a mixed effect regression framework that explicitly accounts for covariate distribution differences across datasets. There is also significant interest in developing harmonization approaches based on deep neural networks (DNNs), such as conditional variational autoencoder (cVAE). However, current DNN approaches do not explicitly account for covariate distribution differences across datasets. Here, we provide mathematical results, suggesting that not accounting for covariates can lead to suboptimal harmonization. We propose two DNN-based covariate-aware harmonization approaches: covariate VAE (coVAE) and DeepResBat. The coVAE approach is a natural extension of cVAE by concatenating covariates and site information with site- and covariate-invariant latent representations. DeepResBat adopts a residual framework inspired by ComBat. DeepResBat first removes the effects of covariates with nonlinear regression trees, followed by eliminating site differences with cVAE. Finally, covariate effects are added back to the harmonized residuals. Using three datasets from three continents with a total of 2787 participants and 10085 anatomical T1 scans, we find that DeepResBat and coVAE outperformed ComBat, CovBat and cVAE in terms of removing dataset differences, while enhancing biological effects of interest. However, coVAE hallucinates spurious associations between anatomical MRI and covariates even when no association exists. Future studies proposing DNN-based harmonization approaches should be aware of this false positive pitfall. Overall, our results suggest that DeepResBat is an effective deep learning alternative to ComBat. Code for DeepResBat can be found here: https://github.com/ThomasYeoLab/CBIG/tree/master/stable_projects/harmonization/An2024_DeepResBat.

## 1 Introduction

There is growing interest in combining MRI data across multiple sites, such as ENGIMA (Thompson et al., 2017) and ABCD (Volkow et al., 2018) studies. These mega-analyses advance neuroimaging research by increasing statistical power (Bethlehem et al., 2022; Marek et al., 2022), enhancing generalizability (He et al., 2022; Lu et al., 2022), and detecting subtle effects (Vogel et al., 2021; Tian et al., 2023). When pooling data across datasets, post-acquisition harmonization is necessary for removing undesirable variabilities across datasets, while preserving relevant biological information. A major source of undesirable cross-dataset heterogeneity is scanner differences across datasets (Magnotta et al., 2012; Chen et al., 2014; Hawco et al., 2018). In addition, the distributions of biological variables (e.g., demographics and clinical diagnosis) may vary across datasets. These biological variables (also referred to as “covariates”) can have a large impact on MRI data (Hua et al., 2010), whose effects should be preserved after harmonization.

A popular approach for harmonizing MRI data is mixed effects models, such as ComBat (Fortin et al., 2017, 2018; Yu et al., 2018). ComBat removes additive and multiplicative site differences while including biological variables as covariates. For example, to perform a mega-analysis using several Alzheimer’s Disease (AD) dementia datasets, the ComBat model might be set up with hippocampal volume as the dependent variable, site as an independent variable, as well as age, sex, and clinical diagnosis as covariates. Additive and multiplicate site effects are removed from the hippocampal volume, while the residual effects of age, sex and clinical diagnoses are retained. Several ComBat variants have been proposed to enhance harmonization performance (Garcia-Dias et al., 2020; Pomponio et al., 2020; Wachinger et al., 2021). However, the simplicity (and elegance) of the ComBat variants limits their ability to eliminate nonlinear site differences spanning the brain regions.

Deep neural networks (DNNs) are promising for eliminating nonlinear site differences distributed across the brain (Dewey et al., 2019; Hu et al., 2023). Variational autoencoder (VAE)-based approaches (Moyer et al., 2020; Russkikh et al., 2020; Zuo et al., 2021; An et al., 2022; Hu et al., 2024) use an encoder to generate site-invariant latent representations from input MRI data. Site information is then concatenated to the latent representations to reconstruct the MRI data via a decoder. Generative adversarial networks (Zhao et al., 2019; Modanwal et al., 2020; Bashyam et al., 2021), normalizing flow (Wang et al., 2021; Beizaee et al., 2023) and federated learning (Dinsdale et al., 2022) have also been explored. However, existing DNN approaches typically overlook the inclusion of covariates, which are explicitly controlled in mixed effects harmonization models (Fortin et al., 2017, 2018; Chen et al., 2022). Since covariate distribution differences are unavoidable across datasets, neglecting covariates during harmonization can inadvertently remove relevant biological information, instead of reducing undesired dataset differences, leading to worse downstream performance. In Section 2.1, we show how a theoretical machine learning result (Tachet et al., 2020) can be used to understand this phenomenon.

In this study, we propose two covariate-aware deep learning harmonization techniques: covariate VAE (coVAE) and deep residual batch effects harmonization (DeepResBat). coVAE extends conditional VAE (cVAE; Moyer et al., 2020) by concatenating covariates and site information with site- and covariate-invariant latent representations. On the other hand, DeepResBat adopts a residual framework inspired by the classical ComBat approach. DeepResBat first removes covariate effects with nonlinear regression trees, followed by eliminating unwanted site differences from the residuals with cVAE. Finally, covariate effects are added back to the harmonized residuals. We found that coVAE hallucinated spurious associations between anatomical MRI and covariates even when no association existed, suggesting that DNN-based harmonization approaches can introduce false positives during harmonization. On the other hand, DeepResBat effectively mitigated this false positive issue.

The contributions of this study are multi-fold. First, we showed theoretically that ignoring covariate differences across datasets can lead to suboptimal harmonization. Second, we introduced a DNN-based harmonization approach DeepResBat that could account for covariate differences across datasets. DeepResBat outperformed ComBat (Fortin et al., 2017), CovBat (Chen et al., 2021) and cVAE (Moyer et al., 2020) across multiple evaluation experiments, including enhancing biological effects of interest, while removing unwanted dataset differences. Third, we demonstrated that DNN-based harmonization approaches could potentially hallucinate relationships between covariates and MRI measurements even when none existed. Therefore, future studies proposing DNN-based harmonization approaches should be aware of this false positive pitfall. Although the current study focused on MRI data, our results are applicable to any field where instrumental harmonization is necessary, e.g., molecular biology (Johnson et al., 2007), geological sciences (Madonna et al., 2022) (Madonna et al., 2022) or agriculture (Leroux et al., 2019).

## 2 Methods

### 2.1 Motivation for accounting for covariates during harmonization

Distribution differences in covariates, such as demographics and clinical diagnoses, across datasets are inevitable. Most deep learning harmonization approaches directly align distributions of latent representations across datasets without explicitly modeling covariate differences (Dewey et al., 2019; Zuo et al., 2021; Beizaee et al., 2023; Liu et al., 2023). Without explicitly accounting for these covariates, the covariates differences can be misinterpreted as undesirable dataset differences and wrongly removed by the harmonization algorithms. Here, we will formalize this phenomenon using a theoretical result from the machine learning literature (Tachet et al., 2020).

More specifically, suppose we have two datasets *S* and *T* with target label *Y* and input data *X*. The goal is to predict *Y* using *X*. Suppose we have feature extractor *g* that takes in *X* as the input with feature representation *Z* as the output. The feature representation *Z* is then entered into classifier *h* to predict the target label. Let ϵ_S_(*h* ∘ *g*) and ϵ_*T*_(*h* ∘ *g*) be the expectation of classification errors when applying *g* followed by *h* to datasets *S* and *T* respectively. Let *Y*_*S*_ and *Y*_*T*_ be the target label distributions in datasets *S* and *T* respectively. Let *Z*_*S*_ and *Z*_*T*_ be the distributions of the feature representation in datasets *S* and *T* respectively. Then, Tachet des Combes and colleagues show that the following inequality is true:

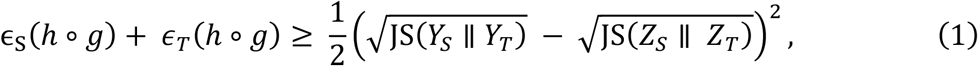

where JS(⋅ ∥ ⋅) is the Jensen-Shannon divergence of two distributions. We note that the lowest possible error bound is zero. Therefore, assuming distributional differences between the target labels of the two datasets (i.e., JS(*Y*_*S*_ ∥ *Y*_*T*_) > 0), then a lower bound of zero can only be achieved if the same distribution differences exist between the feature representations of the two datasets, i.e., JS(*Z*_*S*_ ∥ *Z*_*T*_) = JS(*Y*_*S*_ ∥ *Y*_*T*_). In other words, dataset-invariant representations (i.e., JS(*Z*_*S*_ ∥ *Z*_*T*_) = 0) lead to suboptimal classification performance (greater than zero error bound in Eq. (1)) if there exists distributional differences between the target labels of the two datasets (JS(*Y*_*S*_ ∥ *Y*_*T*_) > 0).

To relate Eq. (1) to harmonization, we can think of *g* as a harmonization procedure (instead of a feature extractor), and *h* as a downstream task to predict covariates *Y* after harmonization. Therefore, if the distributions of covariates *Y* are different across datasets (i.e., JS(*Y*_*S*_ ∥ *Y*_*T*_) > 0), blindly matching the distributions of brain imaging measures across datasets in the harmonization process (i.e., JS(*Z*_*S*_ ∥ *Z*_*T*_) = 0) is suboptimal for the downstream task, i.e., error bound in Eq. (1) is greater than 0.

For example, suppose we would like to harmonize two datasets with different distributions of healthy elderly participants and participants with Alzheimer’s disease (AD) dementia. Then, it is important to account for these distributional differences when harmonizing the datasets. This is typically not an issue for mixed effects harmonization approaches (such as ComBat and CovBat) since covariates are typically explicitly included. However, most deep learning approaches do not account for covariate distribution differences between datasets, which can potentially result in suboptimal downstream task performance.

### 2.2 Datasets and preprocessing

In this study, we proposed DNN models for harmonizing T1 anatomical MRI data. We will test the models using data from separate research initiatives: the Alzheimer’s Disease Neuroimaging Initiative (ADNI) (Jack et al., 2008; Jack et al., 2010), the Australian Imaging, Biomarkers and Lifestyle (AIBL) study (Ellis et al., 2009, 2010; Fowler et al., 2021) and the Singapore Memory Ageing and Cognition Centre (MACC) Harmonization cohort (Hilal et al., 2015; Chong et al., 2017; Hilal et al., 2020). All data collection and analysis procedures were approved by the respective Institutional Review Boards (IRBs), including the National University of Singapore IRB for the analysis presented in this paper. All three datasets encompass a range of modalities collected at multiple timepoints, such as MRI scans, cognition assessments, and clinical diagnoses.

We utilized ADNI1 and ADNI2/Go data from ADNI (Jack et al., 2008; Jack et al., 2010). For ADNI1, the MRI scans were collected from 1.5 and 3T scanners from different vendors (more details in Table S1). For ADNI2/Go, the MRI scans were acquired on 3T scanners. A total of 1,735 participants underwent at least one T1 MRI scan, resulting in 7,955 scans scanned at multiple timepoints (see Table S3 and Figure S1 for details). 68 cortical and 40 subcortical regions of interest (ROI) were defined based on FreeSurfer (Fischl et al., 2002; Desikan et al., 2006). The volumes of the cortical and subcortical ROIs were provided by ADNI using multiple preprocessing steps (http://adni.loni.usc.edu/methods/mri-tool/mri-pre-processing/), followed by the FreeSurfer version 4.3 (ADNI1) and 5.1 (ADNI2/Go) recon-all pipelines, yielding a total of 108 volumetric measures.

In the case of AIBL (Ellis et al., 2009, 2010; Fowler et al., 2021), the MRI scans were collected from 1.5T and 3T Siemens (Avanto, Tim Trio, and Verio) scanners (see Table S2 for more information). There were 495 participants with at least one T1 MRI scan, resulting in 933 MRI scans across multiple timepoints (see Table S3 and Figure S1 for details). The FreeSurfer 6.0 recon-all pipeline was employed to extract the volumes from 108 cortical and subcortical ROIs.

In the case of MACC (Hilal et al., 2015; Chong et al., 2017; Hilal et al., 2020), the MRI scans were collected from a Siemens 3T Tim Trio scanner. There were 557 participants with at least one T1 MRI scan. There were 1197 MRI scans across the different timepoints of the 557 participants (see Table S3 and Figure S1 for details). Similar to AIBL, we utilized the FreeSurfer 6.0 recon-all pipeline to extract the volumes of 108 cortical and subcortical ROIs.

### 2.3 Workflow

To compare different harmonization approaches, we harmonized brain ROI volumes between ADNI and AIBL, as well as ADNI and MACC. Figure 1 illustrates the workflow in this study using AIBL as an example. The procedure was the same for harmonizing ADNI and MACC.

**Figure 1.**
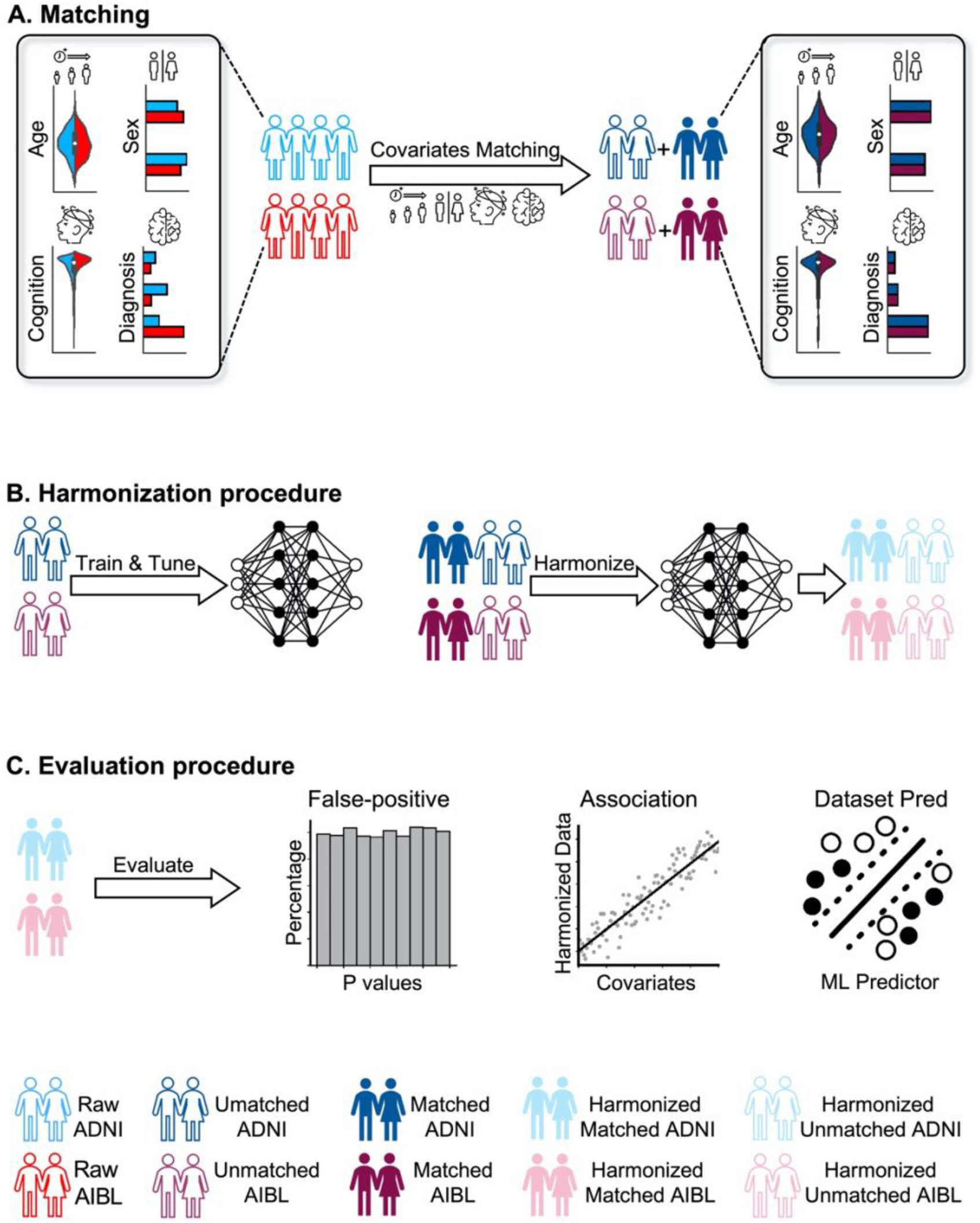
Workflow of study. We illustrate the workflow using ADNI and AIBL. The same procedure was applied to ADNI and MACC. (A) ADNI and AIBL participants were matched based on age, sex, mini mental-state examination (MMSE) and clinical diagnosis. The unmatched participants were used for training and tuning harmonization and evaluation models. The matched participants served as the test set for harmonization evaluation. (B) Left: Train harmonization models with unmatched participants. Right: Apply trained harmonization models on both unmatched and matched participants. (C) Three sets of evaluation experiments were tested on matched harmonized participants: dataset prediction experiment, association analysis and false positive experiment. In evaluations where training was necessary (dataset prediction and false positive experiments), unmatched participants were used as the training and validation sets for the evaluation experiments, while the matched participants were used as the test set.

Following our previous study (An et al., 2022), the Hungarian matching algorithm (Kuhn, 1955) was first applied to match a subset of participants with similar age, sex, Mini Mental State Examination (MMSE, Tombaugh & McIntyre, 1992), and clinical diagnosis distribution between ADNI and AIBL datasets (Figure 1A). The distributions before and after matching are shown in Figure S2. The distributions of remaining unmatched participants are shown in Figure S3. When matching the ADNI and AIBL datasets, we obtained 247 matched participant pairs. When matching the ADNI and MACC datasets, 277 matched participant pairs were obtained. Notably, not all time points had corresponding MMSE and clinical diagnosis information. Therefore, care was taken to ensure that all timepoints in the matched participants had both MMSE and clinical diagnosis. We ensured that each participant’s scans were all categorized as either “matched” or “unmatched” without splitting the participant’s scans across categories. The quality of the matching procedure was assessed through statistical tests, whose p values are reported in Tables S4 to S10. All p values were greater than 0.05.

The resulting unmatched participants were used to train and tune various harmonization approaches (Figure 1B). After model fitting, the trained harmonization models were applied to harmonize matched and unmatched participants. Three evaluation experiments were performed (Figure 1C). First, as a common evaluation practice (Hu et al., 2023), a machine learning model was trained to investigate whether harmonization could effectively reduce dataset differences by predicting which dataset a participant came from (more details in Section 2.8). Second, we evaluated whether harmonization led to stronger associations between harmonized ROI volumes with the covariates of interest (more details in Section 2.9). Finally, an exhaustive false-positive permutation test was carried out, involving 240,000 GPU hours and 360,000 CPU hours. This test aimed to assess whether deep harmonization models might introduce spurious associations (i.e., false positives) between anatomical MRI and randomly permuted covariates when no association exists (more details in Section 2.10). In evaluations where training was necessary (dataset prediction and false positive experiments), unmatched participants were used as the training and validation sets for the evaluation experiments, while the matched participants were used as the test set.

### 2.4 Training, validation and test procedure

As described in section 2.3, the test set for evaluation (Figure 1C) consisted of matched participants. On the other hand, the unmatched participants were utilized for training both harmonization (Figure 1B) and evaluation models (e.g., dataset prediction and false positive experiments in Figure 1C).

In the case of mixed effects models (ComBat and CovBat), there is no hyperparameter, so the data of all unmatched participants were used to fit the models. On the other hand, for the deep harmonization models, we have to tune the hyperparameters. Therefore, we divided the unmatched participants into 10 distinct groups. Notably, all timepoints belonging to a participant were assigned to a single group, thus avoiding any splitting of timepoints of a participant across different groups. For the training and tuning of the deep harmonization models on unmatched participants, a 9-1 train-validation split was employed, with 9 groups used for training and one group used as the validation set for hyperparameter tuning. This process was repeated 10 times, with each group serving as the validation set once. Subsequently, in the harmonization step (Figure 1B), we obtained 10 sets of trained harmonization models. These models were then applied to the unharmonized data, resulting in the generation of 10 harmonized data sets from both unmatched and matched participants.

In the subsequent evaluation step (Figure 1C), the harmonized data of the unmatched participants were used to train and tune the evaluation models for dataset prediction (Section 2.8) and false positive analysis (Section 2.10), following the same 9-1 train-validation split previously described. The harmonized data of the matched participants were designated as the test set to evaluate the harmonization performance. In the case of association analysis (Section 2.9), no evaluation model needed to be trained, so we directly applied general linear models (GLMs) and multivariate analysis of variance (MANOVA) to the harmonized data of the matched participants to obtain association results.

### 2.5 Baseline harmonization models

Here, we considered ComBat (Johnson et al., 2007), CovBat (A. A. Chen et al., 2022), and cVAE (Moyer et al., 2020) as baseline models. We chose ComBat as a baseline since it is the most popular harmonization approach in brain imaging. CovBat is a recent extension of ComBat that seeks to additionally remove site differences in covariance among imaging features, so it is a powerful alternative to ComBat (A. A. Chen et al., 2022). For deep learning baselines, we chose cVAE because of the ease of training cVAE compared with GANs. Furthermore, coVAE is a natural extension of cVAE, so cVAE is a natural baseline to include in our comparisons.

#### 2.5.1 ComBat

ComBat is a mixed effects model that controls for additive and multiplicative site effects (Johnson et al., 2007). Here we utilized the R implementation of the algorithm (https://github.com/Jfortin1/ComBatHarmonization). The ComBat model is as follows:

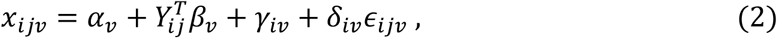

where *i* is the site index, *j* is the participant index, and *v* indexes the brain ROI volumes. *x*_*ijv*_ is the volume of the *v*-th brain ROI of participant *j* from site *i. γ*_*iv*_ is the additive site effect. *δ*_*iv*_ is the multiplicative site effect. *ϵ*_*ijv*_ is the residual error term following a normal distribution with zero mean and variance 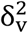. *Y*_*ij*_ is the vector of covariates of participant *j* from site *i*. In this study, we chose age, sex, MMSE, and clinical diagnosis as covariates.

The ComBat parameters *α*_*v*_, *β*_*v*_, *γ*_*iv*_ and *δ*_*iv*_ were estimated for each brain region using the unharmonized ROI volumes of all unmatched participants (Figure 1B). The estimated parameters can then be applied to map a new participant *i* from site *j* to intermediate space with brain regional volume *x*_*ijv*_ and covariates *Y*_*ij*_.

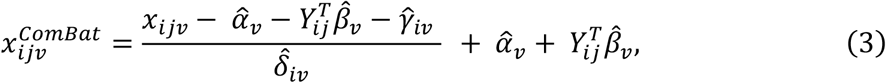

where ^ indicates that the parameter was estimated from the *unmatched unharmonized* ROI volumes from ADNI and AIBL. A separate ComBat model was fitted for harmonizing ADNI and MACC brain regional volumes.

#### 2.5.2 CovBat

CovBat is a mixed effect harmonization model built on top of ComBat to remove site effects in mean, variance, and covariance (Chen et al., 2022). We utilized the authors’ R implementation of the algorithm (https://github.com/andy1764/CovBat_Harmonization). There are four main steps in CovBat harmonization. First, ComBat (Section 2.5.1) is applied to the volume of each brain region. to obtain ComBat adjusted residuals:

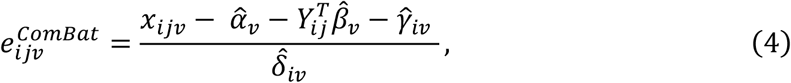

For participant *i* of site *j*, the ComBat-adjusted residuals of all regional volume can be concatenated into a column vector 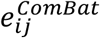. These vectors can in turn be concatenated across all participants of all sites into the matrix *E*^*Combat*^, where the number of columns is equal to the total number of participants and the number of rows is equal to the number of brain regional volume.

Second, principal component analysis (PCA) is applied to *E*^*Combat*^ to obtain *q* principal component (PC) scores and PCs, where *q* is the rank of the matrix *E*^*Combat*^. Therefore, we can write 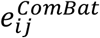 as

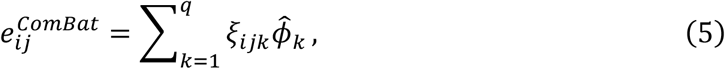

where ξ_*ijk*_ is the *k*-th PC score of participant *i* of site *j*, and 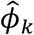 is *k*-th PC.

Third, each of the top *K* PC scores were harmonized using a second round of ComBat, thus yielding 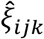. *K* was selected to explain the 95% percentage of variance. The remaining PC scores were not harmonized. Finally, the harmonized brain ROI volumes are projected to intermediate space after model fitting:

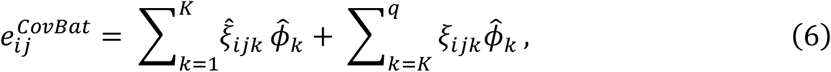

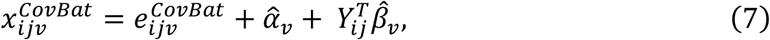

Consistent with ComBat, we chose age, sex, MMSE, and clinical diagnosis as covariates. CovBat was fitted using all unmatched participants.

For a new participant, 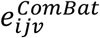 was computed using the ComBat parameters estimated from the unmatched participants. The 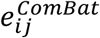 of the new participant was then projected onto the principal components (obtained from the unmatched particiapnts) to obtain principal component scores ξ_*ijk*_. The top *K* scores were then harmonized using the second round of ComBat parameters estimated from the unmatched participants to obtain 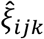. Finally, Equations (6) and (7) were applied to obtain the harmonized ROI volumes of the new participant.

Similar to ComBat, two separate CovBat models were fitted: one for harmonizing ADNI and AIBL brain regional volumes, and one for harmonizing ADNI and MACC brain regional volumes.

#### 2.5.3 cVAE

Moyer et al. (2020) introduced the conditional variational autoencoder (cVAE) model for harmonizing diffusion MRI data. In this paper, we applied the cVAE model to harmonize brain ROI volumes. Figure 2A shows the architecture of the cVAE model. The input brain volumes *x* (*x* denotes brain volumes of all regions: *x*_1,_ *x*_2_ … *x*_*v*_, … *x*_108_) were processed through an encoder deep neural network (DNN) to obtain the latent representation *z*. The one-hot site vector *s* was concatenated with the latent representation *z* and then fed into the decoder DNN, producing the reconstructed brain volumes 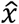. To encourage the independence of the learned representation *z* from the site *s*, the cost function incorporated the mutual information *I*(*z, s*). The resulting loss function could be expressed as follows:

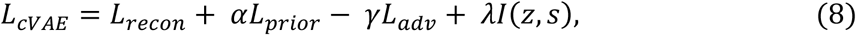

where *L*_*recon*_ was the mean square error (MSE) between *x* and 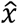, thus encouraging similarity between the harmonized 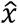 and unharmonized volumes *x*. Additionally, Moyer et al. introduced the term *L*_*adv*_, which was the soft-max cross-entropy loss of an adversarial discriminator aiming to differentiate *x* and 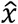, thereby further promoting their similarity. Finally, *L*_*prior*_ was the KL divergence term between representation *z* and a multivariate Gaussian distribution with zero mean and identity covariance matrix (Sohn et al., 2015) to promote regularity and control over the latent space. This prior term will encourage the distributions of latent representations to be aligned across datasets, which can be problematic if there exists covariate differences between datasets (Section 2.1).

**Figure 2.**
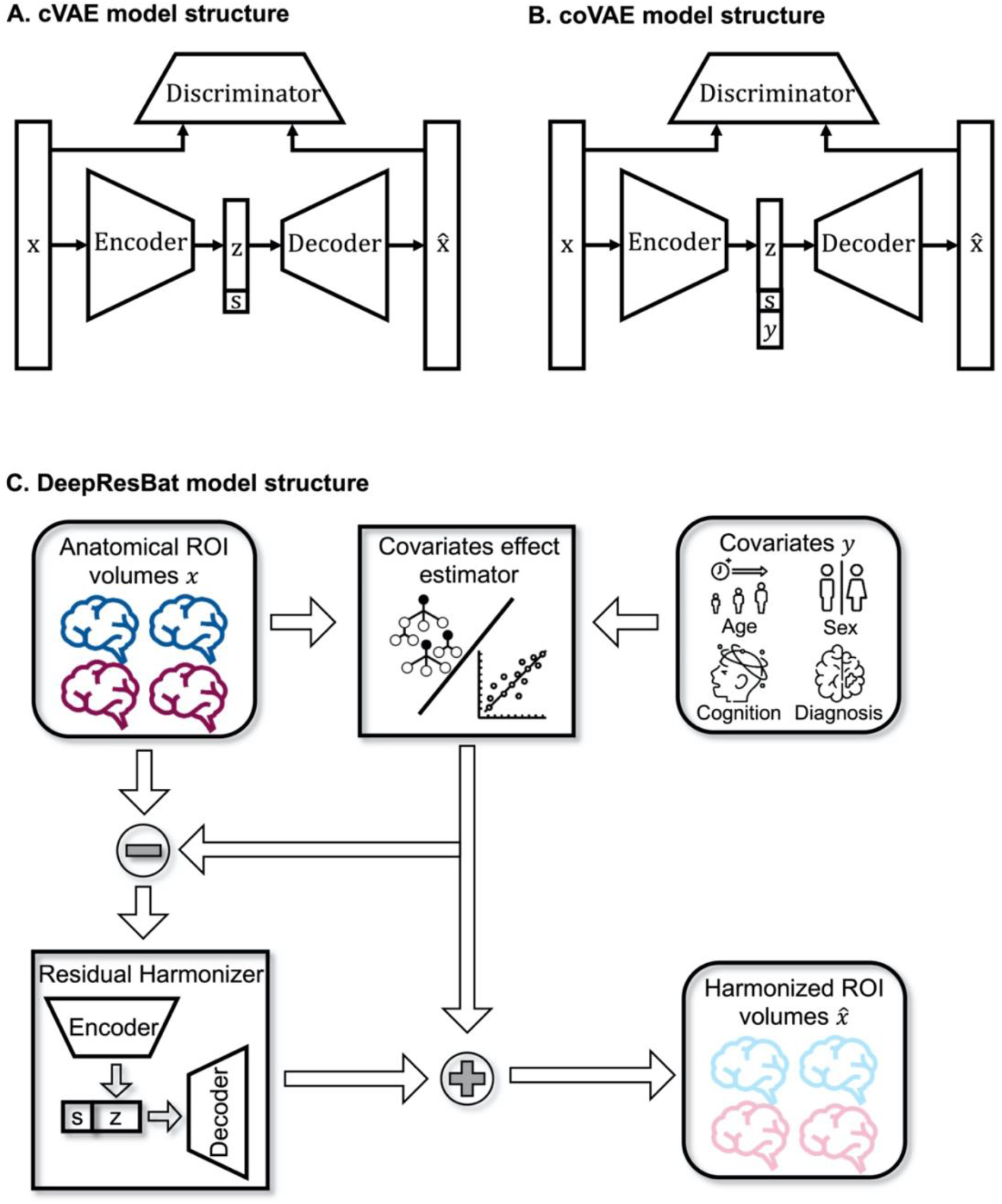
Model structure for cVAE, coVAE, and DeepResBat. (A) Model structure for the cVAE model. Encoder, decoder, and discriminator were all fully connected feedforward DNNs. s was the site we wanted to map the brain volumes to. (B) Model structure for the coVAE model. Site s and covariates y were input into the decoder to preserve covariates effects. Therefore, the main difference between cVAE and coVAE is the inclusion of covariates y. (C) Model structure for the DeepResBat model. The covariates effect estimator was an ensemble of XGBoost and linear models. Once the effects of covariates were removed (subtraction sign), the residual harmonizer was a cVAE model taking covariates-free residuals as input. The covariates effects were then added back to the cVAE output, yielding a final set of harmonized ROI volumes.

The decoder and encoder components of the model were implemented as fully connected feedforward neural networks, where each layer was connected to the subsequent layer. Consistent with Moyer et al., the tanh activation function (Maas et al., 2013) was employed. During training, the variable *s* represented the true site information associated with the input brain volumes *x*. After training, by setting *s* to zero, the input *x* could be mapped to an intermediate space. Training of the model was performed using the data from 90% of the unmatched participants and hyperparameters were tuned using the data from 10% of the unmatched participants as validation set (Section 2.4).

Hyperparameter tuning in the validation set involved optimizing a weighted sum of the reconstruction loss (MSE between *x* and 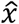) and the accuracy of participant-level dataset prediction: ½ MSE + Dataset Prediction Accuracy. The reconstruction loss (MSE) was halved to ensure comparability with the dataset prediction accuracy. Dataset prediction accuracy was determined by training an XGBoost classifier on the training set and evaluating it on the validation set. To identify the best set of hyperparameters, the HORD algorithm (Eriksson et al., 2020; Regis & Shoemaker, 2013; Ilievski et al., 2017) was employed using the validation set (Table 1). The trained deep neural network (DNN) was then utilized for subsequent analyses after 1000 epochs of training.

**Table 1.** Hyperparameters search ranges for cVAE on the validation set. We note that a learning rate decay strategy was utilized. After K training epochs (where K = learning rate step), the learning rate was reduced by a factor of 10.

| Hyperparameter | Search Range |
| --- | --- |
| Initial learning rate | 1e-2 – 1e-1 |
| Learning rate step | 10 - 999 |
| Dropout rate | 0 – 0.5 |
| $\alpha$ | 0.01 - 1 |
| $\gamma$ | 0.01 - 10 |
| $\lambda$ | 0.01 - 1 |
| Nodes for each layer | 32 - 512 |
| Number of layers | 2 - 4 |
| Node for $z$ | 32 - 512 |

### 2.6 coVAE

As highlighted in Section 2.1, accounting for covariates is essential for effective harmonization. Therefore, we extended the cVAE model to incorporate covariates as input, resulting in the covariate-VAE (coVAE) model, as shown in Figure 2B. More specifically, we concatenated site *s* and covariates *Y* to obtain [*s, Y*]. The loss function was the same as cVAE (Eq. (8)) except that the mutual information loss term in Eq. (8) was modified to become *I*(*z*, [*s, Y*]):

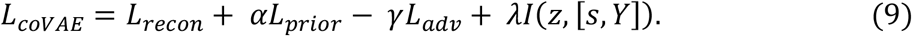

Therefore, instead of minimizing the mutual information between *z* and s, we minimize the mutual information between *z* and [*s, Y*]. For the reconstruction, we concatenated latent representation *z* with site *s* and covariates *Y* as input to the decoder network.

Recall that the *L*_*prior*_ term was the KL divergence term between representation *z* and a multivariate Gaussian distribution with zero mean and identity covariance matrix. This prior term therefore implicitly encouraged the alignment of the latent representations *z* between datasets. In the case of cVAE, the latent representation *z* would contain covariate information. Therefore, cVAE would force the alignment of latent representations *z* even if the covariate distributions were different across datasets, which would be suboptimal (see Section 2.1). By contrast, because coVAE seek to minimize mutual information between *z* and [*s, Y*], so the latent representation *z* would theoretically not contain covariate information. In this scenario, aligning the covariate-free and site-free latent representation *z* would make sense.

Consistent with ComBat (Section 2.5.1) and CovBat (Section 2.5.2), we chose age, sex, MMSE, and clinical diagnosis as covariates. The categorical covariates sex and clinical diagnosis were one-hot encoded for coVAE. Training of coVAE was performed using the data from 90% of the unmatched participants and hyperparameters were tuned using the data from 10% of the unmatched participants as validation set (Section 2.4). We used the HORD algorithm to search within the hyper-parameter ranges specified in Table 1 based on the validation set. Brain ROI volumes were mapped to intermediate space for subsequent analyses after 1000 training epochs.

Although coVAE appeared to be an intuitive straightforward extension of cVAE, as will be shown in the false positive analysis (Section 3.3), coVAE suffered from significant false positive rates. Therefore, in the next section, we proposed a second harmonization approach that could account for covariate distribution differences across datasets.

### 2.7 DeepResBat

Figure 2C illustrates our proposed DeepResBat approach. To motivate DeepResBat, we can write ComBat’s Eq. (2) into a more general form:

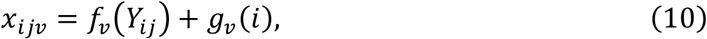

where *i* is the site index, *j* is the participant index, and *v* indexes the brain ROI volumes. *x*_*ijv*_ is the *v*-th brain volume of participant *j* from site *i. Y*_*ij*_ are the covariates of participant *j* from site *i*. In ComBat, *f*_*v*_ is linear with 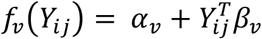, while *g*_*v*_(*i*) = *γ*_*iv*_ + *δ*_*iv*_*ϵ*_*ijv*_ accounts for both additive and multiplicative site effects.

To improve on ComBat, DeepResBat utilized nonlinear functions for *f* and *g*. There are three stages for DeepResBat (Figure 2C). We first estimated the covariate effects *f* using a nonlinear regression approach (Section 2.7.1). The covariate-free residuals from the first stage (*x* − *f*) were then harmonized using a generic deep learning approach, instantiated as cVAE in the current study (Section 2.7.2). The covariate effects from the first stage were then added back to the harmonized brain volumes from the second stage (Section 2.7.3).

Consistent with ComBat, we chose age, sex, MMSE, and clinical diagnosis as covariates. Training of DeepResBat was performed using the data from 90% of the unmatched participants and hyperparameters were tuned using the data from 10% of the unmatched participants as validation set (Section 2.4). We used the HORD algorithm to search within the hyper-parameter ranges specified in Table 1 based on the validation set. Brain ROI volumes were mapped to intermediate space for subsequent analyses after 1000 training epochs.

#### 2.7.1 Covariates effects estimation

To estimate covariate effects, we first regressed out the linear effects of site for each brain ROI volume by fitting the following model,

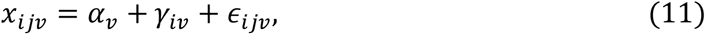

where *x*_*ijv*_ is the *v*-th brain ROI volume for participant *i* of site *j, α*_*v*_ is the intercept term, *γ*_*iv*_ is the additive sites effect, and *ϵ*_*ijv*_ is the residual error term. The residual brain ROI volume 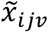 without linear site effects is obtained by 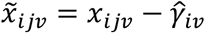.

Before removing covariate effects, we first checked whether each covariate is actually related to any ROI brain volume in order to avoid the false positive issues exhibited by coVAE. This check is performed in two stages. First, for each covariate, an XGBoost model (T. Chen & Guestrin, 2016) was trained to predict the covariate using all residual brain ROI volumes 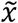 obtained from the previous step (Eq. (11)). XGBoost was chosen due to its efficacy with unstructured or tabular data (Grinsztajn et al., 2022; Shwartz-Ziv & Armon, 2022) and its simplicity, allowing for fast training. For each covariate, the XGBoost model was trained using the training set (90% of unmatched participants) and hyperparameters were tuned using the validation set (10% of unmatched participants). We randomly sampled 50% of participants from the validation set and computed the correlation between the prediction and ground truth covariate. Pearson’s correlation and Spearman’s correlation were used for continuous and discrete covariates respectively. This sampling procedure was repeated 100 times. If the p values of the correlations were less than 0.05 for more than 95% of the repetitions, then we retained the covariate for the next stage.

In the second stage, for each brain volume 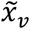, an XGBoost model was trained to predict the brain volume using all survived covariates 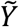. Once again, the training used the training set (90% of unmatched participants) and hyperparameters were tuned using the validation set (10% of unmatched participants). To ensure, we were not overfitting the covariate estimator, we again sampled 50% of participants from the validation set and computed the correlation between the prediction and ground truth covariate. Pearson’s correlation was used for evaluating brain ROI volumes’ predictions. This sampling procedure was repeated 100 times. If the p values of the correlations were less than 0.05 for more than 95% of the repetitions, we retained the XGBoost model. If not, we fitted a linear model instead.

Therefore, regardless of whether we ended up using a linear model or XGBoost, we obtained the covariates effect estimator 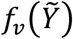 for each brain region *v*. The estimated covariates effects could then be subtracted from the original brain ROI volume, yielding covariate-free residuals:

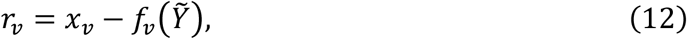

The residuals *r*_*v*_ were presumably free from covariate effects, but retained unwanted variations from each dataset, which could be removed with a generic deep learning based harmonization approach in the next stage (Section 2.7.2).

#### 2.7.2 Covariate-free residuals harmonization

In the second stage of DeepResBat, the covariate-free residuals *r*_*v*_ were jointly fed into a deep learning based harmonization model *g*(⋅) for further harmonization. In the current study, we chose the cVAE model, although any deep learning harmonization model could be used. Similar to the cVAE baseline (Section 2.5.3), the cVAE was trained using the training set (90% of unmatched participants) and hyperparameters (Table 1) were tuned using the validation set (10% of unmatched participants) with the HORD algorithm.

Following training, the covariates-free residuals were mapped to an intermediate space:

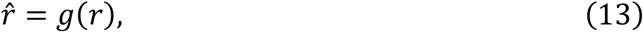

where *r* was the covariate-free residuals and 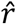 was the harmonized residual brain volumes.

#### 2.7.3 DeepResBat harmonization

The final harmonized brain ROI volumes were then obtained by adding the estimated covariates effect from stage 1 and harmonized residual from stage 2 for each brain ROI volume:

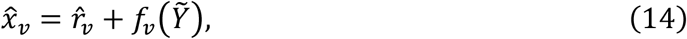

where 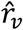 was the harmonized residuals (Section 2.7.2) and 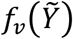 was the estimated covariate effects (2.7.1).

### 2.8 Dataset prediction model

As an evaluation metric, we employed XGBoost to predict the source dataset of the harmonized brain volumes (Figure 1C). The inputs to the XGBoost model were the brain volumes normalized by each participant’s total intracranial volume (ICV). Due to the 10-fold cross-validation procedure described in Section 2.4, recall that the unmatched participants were divided into 10 groups of training and validation sets. Therefore, in the case of cVAE, coVAE and DeepResBat, there were 10 harmonization models and 10 sets of harmonized data for each participant. In the case of ComBat and CovBat, the models were fitted on all unmatched participants, so there was only one set of harmonized data for each participant.

For each group of training and validation sets, an XGBoost classifier was trained using the training set and a grid search was conducted on the validation set to identify the optimal hyperparameters. To evaluate performance, the 10 XGBoost classifiers were used to predict the source dataset of the harmonized MRI volumes of the matched participants.

The prediction accuracy was calculated by averaging the results across all time points of each participant and the 10 classifiers before further averaging across participants. To evaluate the harmonization quality between ADNI and AIBL, this evaluation procedure was applied to the ADNI and AIBL participants. The same procedure was applied to evaluate the harmonization quality between ADNI and MACC datasets. Lower prediction accuracies indicated that greater dataset differences were removed, suggesting better harmonization quality.

### 2.9 Association analysis

Lower dataset prediction accuracies (Section 2.8) indicate greater dataset differences were removed, but the removed dataset differences might contain important biological information, which should not be removed. Therefore, association analyses (using the python package *statsmodels*) were conducted to evaluate the preservation of relevant biological information (Figure 1C) after harmonization.

The variables of interest were age, sex, clinical diagnosis, and MMSE. Univariate and multivariate association analyses were performed on matched participants using unharmonized or harmonized regional brain volumes from different approaches. As mentioned in previous sections, brain ROI volumes were harmonized by mapping to intermediate space.

Of the 108 brain regions, 87 regions were grey matter ROIs. Grey matter volumes are known to be correlated with age, sex, cognition, and AD dementia (Hutton et al., 2009; Hua et al., 2010; Blessed et al., 2018; van de Mortel et al., 2021), so the association analyses were focused on the 87 grey matter ROIs.

We again remind the reader that in the case of cVAE, coVAE and DeepResBat, there were 10 harmonization models and 10 sets of harmonized data for each participant. Therefore, the 10 sets of harmonized data were averaged for each matched participant before the association analysis was performed. In the case of ComBat and CovBat, there was only one set of harmonized data for each matched participant, so no averaging was necessary.

#### 2.9.1 Univariate association analysis

For each grey matter brain ROI volume and each harmonization approach, GLM models were fitted to evaluate the association between brain ROI volume and covariates. We fitted GLM separately for clinical diagnosis and MMSE to avoid cofounding:

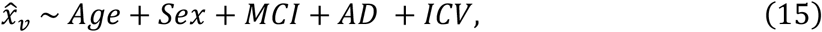

and

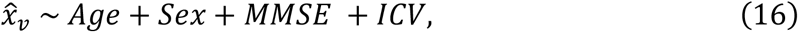

where 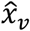 was the *v*-th harmonized brain ROI volume. Z statistics for GLM’s betas were utilized as evaluation metrics.

#### 2.9.2 Multivariate association analysis

Multivariate analysis of variance (MANOVA) was applied to evaluate the multivariate association between grey matter ROIs and covariates. MANOVA was run separately for clinical diagnosis and MMSE:

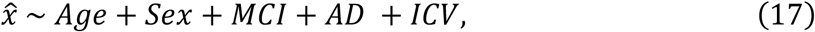

and

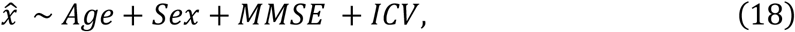

where 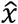 referred to all 87 harmonized grey matter brain ROI volumes. Effect size was measured with Pillai’s Trace (O’Brien & Kaiser, 1985) where a larger Pillai’s Trace value indicated a stronger association.

### 2.10 False positive analysis

To evaluate whether the harmonization approaches will hallucinate associations between covariates and harmonized ROI volumes when none exists, we performed a permutation analysis to evaluate false positive rates. For example, if we permuted age across participants, then the resulting harmonized brain ROI volumes should not associate with the permuted age.

More specifically, when harmonizing ADNI and MACC, for each permutation, we randomly shuffled four covariates (age, sex, MMSE, and clinical diagnosis) together across (1) unmatched participants in the training set, (2) unmatched participants in the validation set and (3) matched participants. Harmonization models were then trained to harmonize brain ROI volumes based on the randomly shuffled covariates. As stated in Section 2.4, unmatched participants were used for training and tuning the harmonization models, and GLMs were performed in the matched harmonized participants.

We expect the association between the randomly permuted covariates and harmonized brain ROI volumes to not exist. Therefore, we ran GLMs to validate our assumption via association analysis. The GLMs were run for each harmonized brain ROI volume with randomly shuffled covariates on matched participants. We considered 87 grey matter ROIs (see Section 2.9.1) to run the GLM. A diagnosis GLM 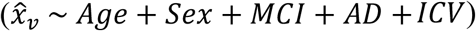 and a cognition 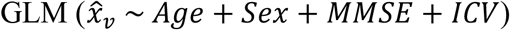 were fitted separately. Then for each permutation and each harmonized brain ROI volume, we obtained p values from the GLM corresponding to each covariate.

This permutation procedure was repeated 1000 times. For each brain ROI volume harmonized by each harmonization model, we calculate the percentage of significant p values below 0.05 across the 1000 p values from the 1000 permutations. The expected percentage across all grey matter ROIs is 5%, with a confidence interval (CI) of 3.65% to 6.35% based on the normal approximation of the Binomial 95% CI (Eklund et al., 2016). Percentage higher than 6.35% indicated that there were false positives. The same procedure was repeated for ADNI and AIBL datasets.

### 2.11 Further analyses

We performed three additional analyses to explore the practical considerations of applying the proposed DeepResBat algorithm.

#### 2.11.1 Sample size requirements for DeepResBat

To investigate the sample size required for DeepResBat, we repeated the dataset prediction and MANOVA analyses (Figures 1), but the sample size used to fit the harmonization models was varied by randomly sampling 50, 60, 70, 80, 90, 100, 110, 120, 130, 140, 150 MRI scans per dataset from the unmatched participants. Similar to the main analyses, 90% of MRI scans were used to train DeepResBat, while the remaining 10% of MRI scans were used as the validation set. For ComBat, all sampled MRI scans were used to fit the model. The fitted harmonization models were applied to matched unharmonized participants for evaluation. We repeated this procedure 50 times. The procedure was performed for ADNI-AIBL and ADNI-MACC cohorts separately.

#### 2.11.2 Harmonization across many sites

Our main analyses involved harmonizing pairs of datasets. In modern mega-analyses, harmonization might need to be performed across many sites. DeepResBat can be easily extended to many sites by extending the cVAE site variable *s* to be a one-hot site vector. To evaluate the harmonization of many sites, we performed a pseudo-site harmonization experiment. In particular, we defined a pseudo-site as a batch of MRI scans acquired on same scanner model and preprocessed with the same FreeSurfer version.

For simplicity, we only considered the first MRI scan of each participant. We then removed pseudo-sites with less than 20 samples, yielding 17 pseudo-sites from the ADNI dataset, 3 pseudo-sites from the AIBL dataset, and 1 pseudo-site from the MACC dataset. We performed a 5-fold cross-validation scheme, where each fold has around 20% of MRI scans from each pseudo-site. For DeepResBat, we used 3 folds as the training set, 1 fold as the validation set, and 1 fold as the test set. For fair comparison, we used 4 folds to fit the ComBat and CovBat models given that these two approaches do not have hyperparameters to tune. The sample characteristics of the 21 pseudo sites are shown in Figure 3. We chose site prediction accuracy and effect size measured by MANOVA as our evaluation metrics.

**Figure 3.**
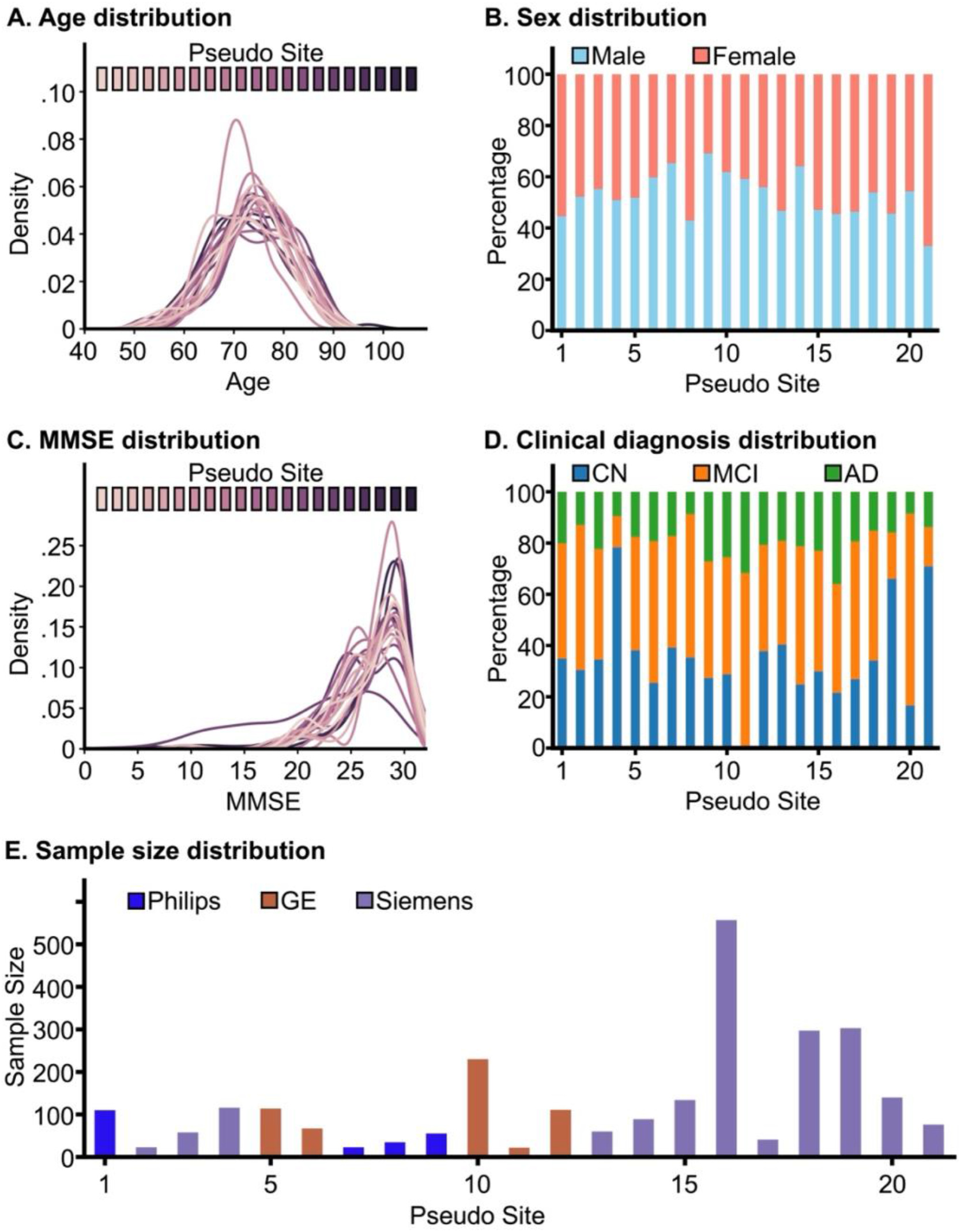
Sample characteristics of 21 pseudo sites. (A) Age distribution for each pseudo site; (B) Sex distribution for each pseudo site; (C) MMSE distribution for each pseudo site; (D) Clinical diagnosis distribution for each pseudo site; (D) Sample size distribution for each pseudo site, colored by MRI vendors.

#### 2.11.3 Harmonization performance based on imbalanced test set

Our main experiments were tested on participants with matched covariates (Figure 1). We further evaluated DeepResBat by subsampling matched participants to yield test sets with highly imbalanced covariates.

Let us consider the case of harmonizing ADNI and AIBL datasets. To obtain an age-imbalanced test set, we considered the oldest 30% matched unharmonized ADNI participants and youngest 30% matched unharmonized AIBL participants. To obtain a sex-imbalanced test set, we randomly sampled 90% male and 10% female from the matched ADNI participants, as well as randomly sampled 10% male and 90% female from the matched AIBL participants. To obtain an MMSE-imbalanced test set, we considered the 30% of matched ADNI participants with highest MMSE scores and 30% of matched AIBL participants with the lowest MMSE scores. To obtain a diagnosis-imbalanced set, we only considered matched ADNI participants with AD or MCI and matched AIBL participants with CN or MCI. The same subsampling procedure was repeated for matched ADNI and MACC participants.

For the evaluation metric, we considered effect size of MANOVA (similar to Section 2.9.2). We note that dataset prediction accuracy is not a useful metric here because of huge covariate differences between the two datasets we are seeking to harmonize. Therefore, a good harmonization algorithm that retains covariate effects (as desired) should exhibit a large dataset prediction accuracy.

### 2.12 Deep neural network implementation

The DNNs developed in this paper were implemented using PyTorch (Paszke et al., 2017) and executed on NVIDIA RTX 3090 GPUs with CUDA 11.0. The DNNs were optimized using the Adam optimizer (Kingma & Ba, 2017) with the default settings provided by PyTorch.

### 2.13 Statistical tests

To assess distribution differences in age between matched participants of AIBL and ADNI (as well as MACC and ADNI), two-sided two-sample t-tests were employed. For sex and clinical diagnoses, chi-squared tests were utilized to examine any significant distinctions. For MMSE, the Kolmogorov-Smirnov test was performed.

In the case of dataset prediction, for cVAE, coVAE and DeepResBat, the prediction performance was averaged over all time points of each participant and then across the 10 sets of models, resulting in a single prediction performance value for each participant. In the case of ComBat and CovBat, there is only one set of models, so the prediction performance was averaged over all time points of each participant, which again yielded a single prediction performance value for each participant. Overall, this process yielded a vector of prediction performance for each dataset and harmonization approach, with each element corresponding to a particular participant. To compare the dataset prediction performance between the two harmonization approaches, a permutation test with 10,000 permutations was conducted. Each permutation involved randomly exchanging the entries between the performance vectors of the two approaches. A more detailed illustration of this permutation procedure can be found in Figure S4.

Multiple comparisons were corrected with a false discovery rate (FDR) of q < 0.05.

### 2.14 Data and code availability

Code for the various harmonization algorithms can be found here (https://github.com/ThomasYeoLab/CBIG/tree/master/stable_projects/harmonization/An2024_DeepResBat). Two co-authors (CZ and NW) reviewed the code before merging it into the GitHub repository to reduce the chance of coding errors.

The ADNI and the AIBL datasets can be accessed via the Image & Data Archive (https://ida.loni.usc.edu/). The MACC dataset can be obtained via a data-transfer agreement with the MACC (http://www.macc.sg/).

## 3 Results

### 3.1 DNN models removed more dataset differences than classical mixed effect models

Dataset prediction accuracies of matched participants are shown in Figure 4. Lower prediction accuracies indicated that greater dataset differences were removed, suggesting better harmonization quality.

**Figure 4.**
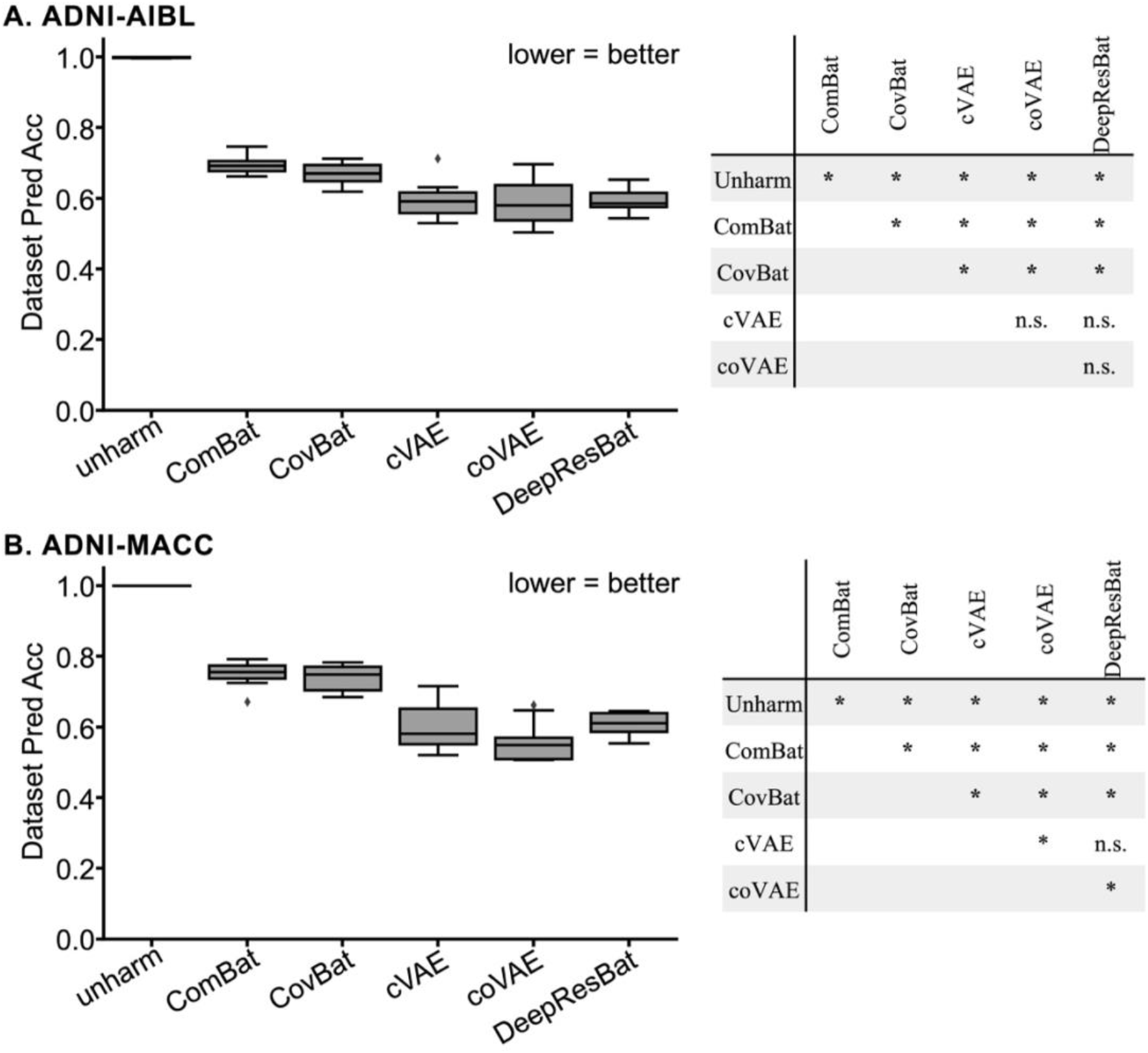
Dataset prediction accuracies. (A) Left: Dataset prediction accuracies for matched ADNI and AIBL participants across 10 folds. Right: p values of differences between different approaches. “*” indicates statistical significance after surviving FDR correction (q < 0.05). “n.s.” indicates not significant. (B) Same as (A) but for matched ADNI and MACC participants. All p values are reported in Tables 2 and 3.

Figure 4A shows the dataset prediction performance for matched ADNI and AIBL participants. Without harmonization, an XGBoost classifier achieved 100% accuracy in identifying which dataset a participant’s data came from. After applying mixed effect harmonization approaches (ComBat and CovBat), the prediction accuracy significantly dropped to 0.695 ± 0.376 (mean ± std) for ComBat, and 0.670 ± 0.383 for CovBat, indicating a substantial reduction in dataset differences. Deep learning approaches showed improved dataset difference removal, with performance of 0.595 ± 0.381 for cVAE and 0.592 ± 0.368 for coVAE. Our proposed DeepResBat achieved an accuracy of 0.594 ± 0.375, which was not statistically different from the deep learning baselines (Table 2). Notably, all deep learning approaches exhibited significantly lower dataset prediction accuracies than mixed effect approaches, demonstrating the potential of deep learning for data harmonization. However, the dataset prediction accuracies of all deep learning approaches remained better than chance (p = 1e-4), indicating residual dataset differences.

**Table 2.** Dataset prediction accuracies with p values of differences between different approaches for matched ADNI and AIBL participants. Statistically significant p values after FDR (q < 0.05) corrections are bolded.

| Dataset Prediction Accuracies<br>(mean ± std) | p values | | | | | |
| --- | --- | --- | --- | --- | --- | --- |
|  | Unharm | ComBat | CovBat | cVAE | coVAE | DeepResBat |
| Unharmonized (1.000 ± 0.020) | | **1e-4** | **1e-4** | **1e-4** | **1e-4** | **1e-4** |
| ComBat (0.695 ± 0.376) | | | **8e-4** | **1e-4** | **1e-4** | **1e-4** |
| CovBat (0.670 ± 0.383) | | | | **1e-4** | **1e-4** | **1e-4** |
| cVAE (0.595 ± 0.381) | | | | | 0.5945 | 0.8931 |
| coVAE (0.592 ± 0.368) | | | | | | 0.8303 |
| DeepResBat (0.594 ± 0.375) | | | | | | |

**Table 3.** Dataset prediction accuracies with p values of differences between different approaches for matched ADNI and MACC participants. Statistically significant p values after FDR (q < 0.05) corrections are bolded.

| Dataset Prediction Accuracies<br>(mean ± std) | p values | | | | | |
| --- | --- | --- | --- | --- | --- | --- |
|  | Unharm | ComBat | CovBat | cVAE | coVAE | DeepResBat |
| Unharmonized (1.000 ± 1e-16) | | **1e-4** | **1e-4** | **1e-4** | **1e-4** | **1e-4** |
| ComBat (0.752 ± 0.361) | | | **0.0095** | **1e-4** | **1e-4** | **1e-4** |
| CovBat (0.740 ± 0.370) | | | | **1e-4** | **1e-4** | **1e-4** |
| cVAE (0.603 ± 0.391) | | | | | **1e-4** | 0.7258 |
| coVAE (0.558 ± 0.418) | | | | | | **2e-4** |
| DeepResBat (0.608 ± 0.381) | | | | | | |

Similar outcomes were observed for matched ADNI and MACC participants (Figure 4B). Without harmonization, the XGBoost classifier accurately predicted source datasets with 100% accuracy. Mixed effect approaches, including ComBat and CovBat, reduced dataset differences to some extent, yielding accuracies of 0.752 ± 0.361 for ComBat and 0.740 ± 0.370 for CovBat. All deep learning approaches (cVAE: 0.603 ± 0.391, coVAE: 0.558 ± 0.418, DeepResBat: 0.608 ± 0.381) exhibited more effective removal of dataset differences than mixed effect approaches (Table 3). Our proposed DeepResBat achieved similar accuracy to cVAE with no statistical difference, but performed worse than coVAE, indicating room for improvement. However, coVAE introduced significant false positives (as will be shown in Section 3.3) and is therefore not an acceptable approach. Finally, the dataset prediction accuracies of all deep learning approaches remained better than chance (p = 1e-4), indicating the presence of residual dataset differences.

We note the relatively high standard deviation (across participants) in dataset prediction accuracies of more than 0.3 across all approaches. Figure S5 shows the histogram of participant-level dataset prediction accuracies for DeepResBat. We found that the accuracies were bimodal with peaks at 0 and 1. One reason is that most matched participants only have one timepoint and the predictions were relatively consistent across the 10 dataset prediction classifiers. Therefore, the predictions were often consistently wrong (or right) for a given participant. The bimodal distribution led to a relatively high variance.

### 3.2 DeepResBat enhanced associations between harmonized brain volumes and covariates

To ensure that relevant biological information was retained in the harmonization process, we performed association analyses between harmonized grey matter volumes and covariates.

#### 3.2.1 DeepResBat outperformed baselines for univariate analysis

Figures 5 and 6 show the results of the univariate GLM association analysis involving clinical diagnosis in the ADNI-AIBL and ADNI-MACC matched participants respectively. Each dot in the plots represented a different brain region, so there are 87 dots in total. For age, MCI and AD dementia, more negative z values indicated greater atrophy due to aging and AD progression. Conversely, a lower MMSE indicates worse cognition, so more positive z values indicated greater atrophy related to worse cognition. For sex, the absolute z statistics were compared because there was no a priori expectation of positive or negative values, so a larger magnitude indicating a larger effect size. Overall, red dots indicate better performance by DeepResBat, while blue dots indicate worse performance.

**Figure 5.**
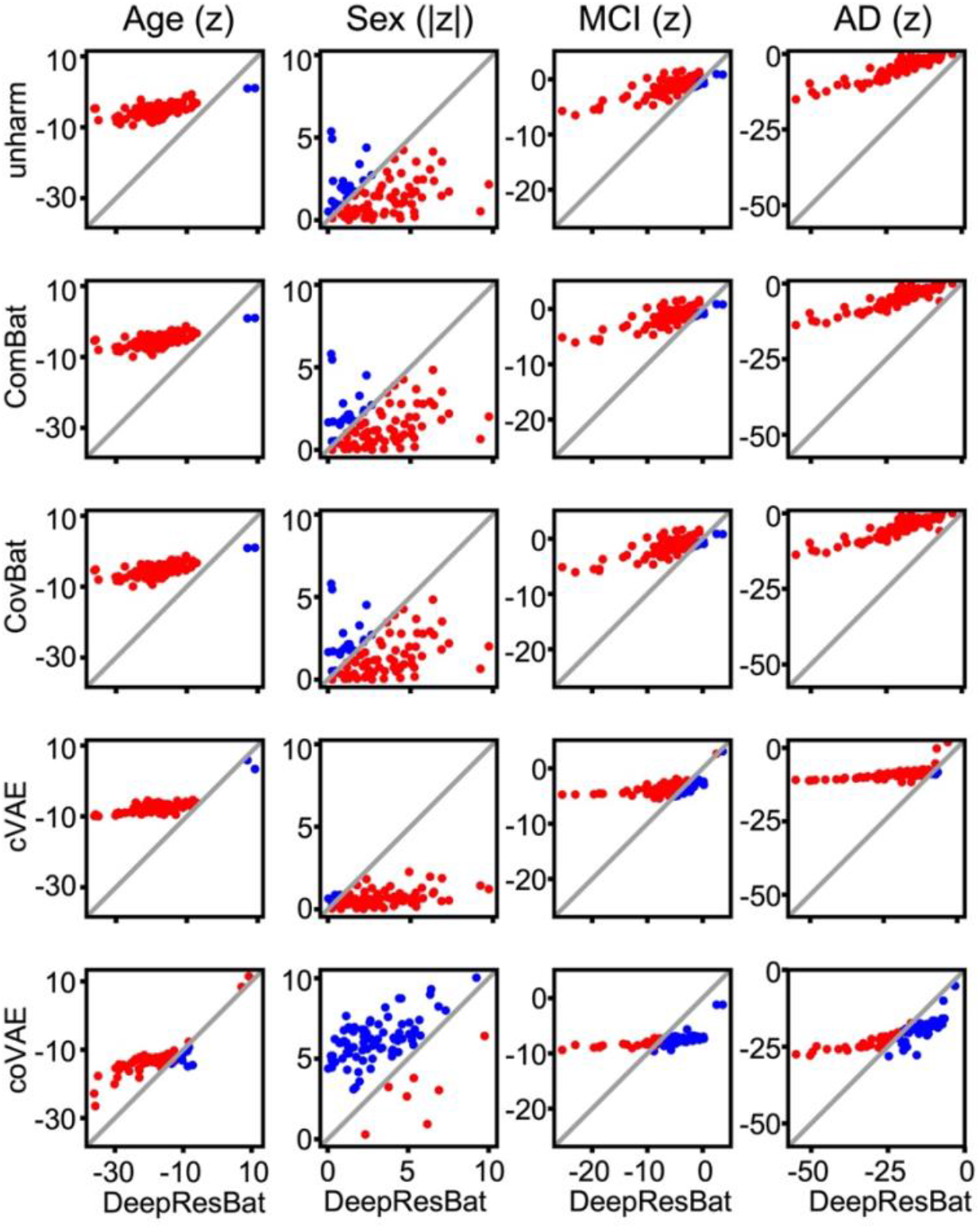
Comparison of z statistics from GLM involving clinical diagnosis for DeepResBat and baselines on matched ADNI and AIBL participants. Each row compares DeepResBat and one baseline approach: no harmonization (row 1), ComBat (row 2), CovBat (row 3), cVAE (row 4) and coVAE (row 5). Each column represents one covariate: age (column 1), sex (column 2), MCI (column 3) and AD dementia (column 4). Each subplot compares z statistics of DeepResBat against another baseline for a given covariate across 87 grey matter ROIs. Each dot represents one grey matter ROI. Red dots indicate better performance by DeepResBat. Blue dots indicate worse performance by DeepResBat.

**Figure 6.**
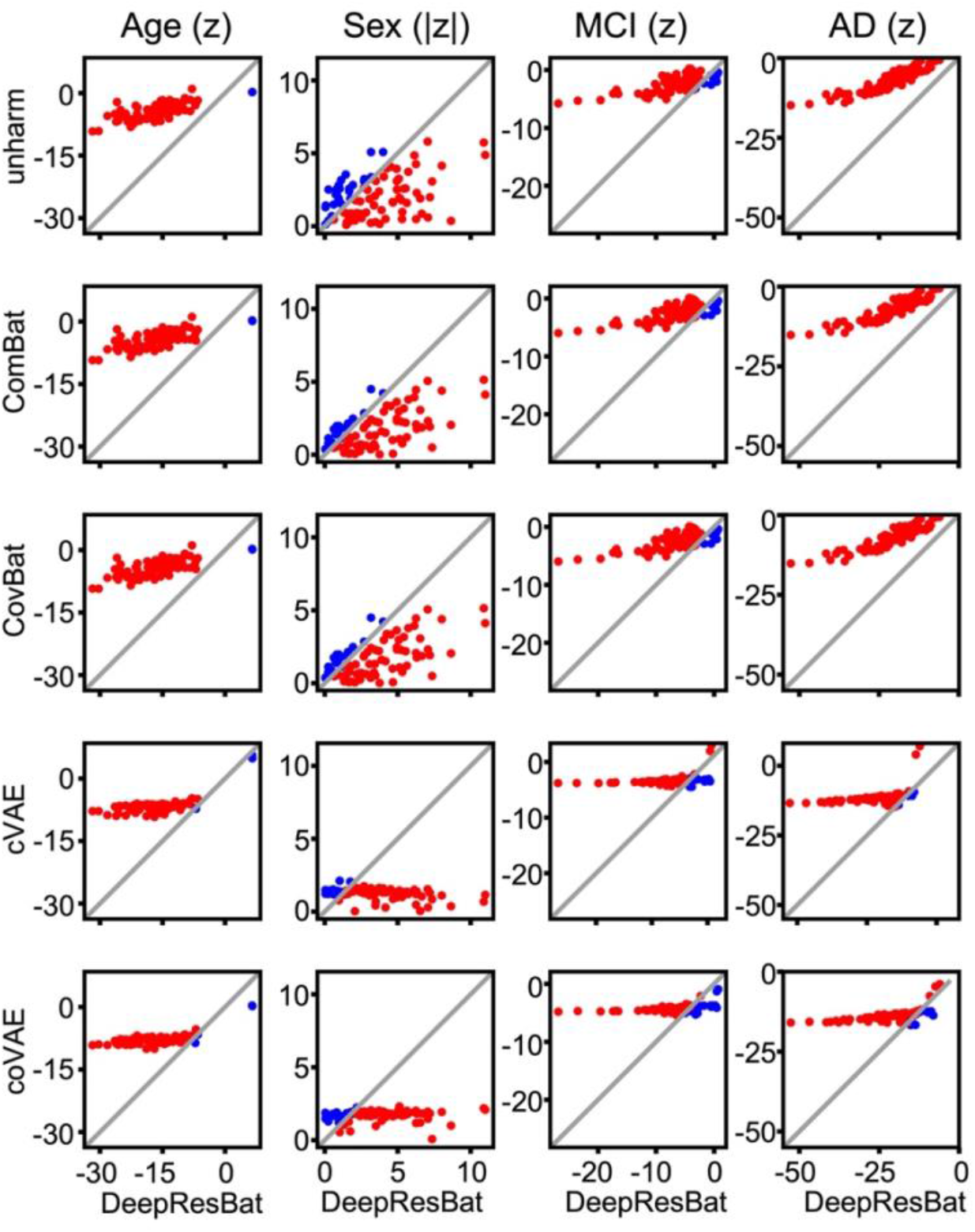
Comparison of z statistics from GLM involving clinical diagnosis for DeepResBat and baselines on matched ADNI and MACC participants. Each row compares DeepResBat and one baseline approach: no harmonization (row 1), ComBat (row 2), CovBat (row 3), cVAE (row 4) and coVAE (row 5). Each column represents one covariate: age (column 1), sex (column 2), MCI (column 3) and AD dementia (column 4). Each subplot compares z statistics of DeepResBat against another baseline for a given covariate across 87 grey matter ROIs. Each dot represents one grey matter ROI. Red dots indicate better performance by DeepResBat. Blue dots indicate worse performance by DeepResBat.

In the case of matched ADNI-AIBL participants, DeepResBat yielded stronger associations between brain volumes and all covariates with respect to no harmonization, ComBat, CovBat and cVAE (Figure 5 rows 1 to 4). Compared to coVAE, DeepResBat yielded weaker absolute z statistics for sex, similar z statistics for MCI and AD, and better z statistics for age (Figure 5 row 5). However, coVAE introduced false positives (Section 3.3) and is therefore not an acceptable approach. In the case of matched ADNI-MACC participants, DeepResBat yielded stronger associations between brain volumes and all covariates with respect to no harmonization and all other baselines (Figure 6).

Similar conclusions were obtained for the univariate GLM association analyses involving MMSE (Figures S6 and S7).

#### 3.2.2 DeepResBat outperformed baselines for multivariate analysis

Figure 7 shows the results of the multivariate MANOVA association analysis involving clinical diagnosis in the ADNI-AIBL and ADNI-MACC matched participants. A higher bar indicates better performance.

**Figure 7.**
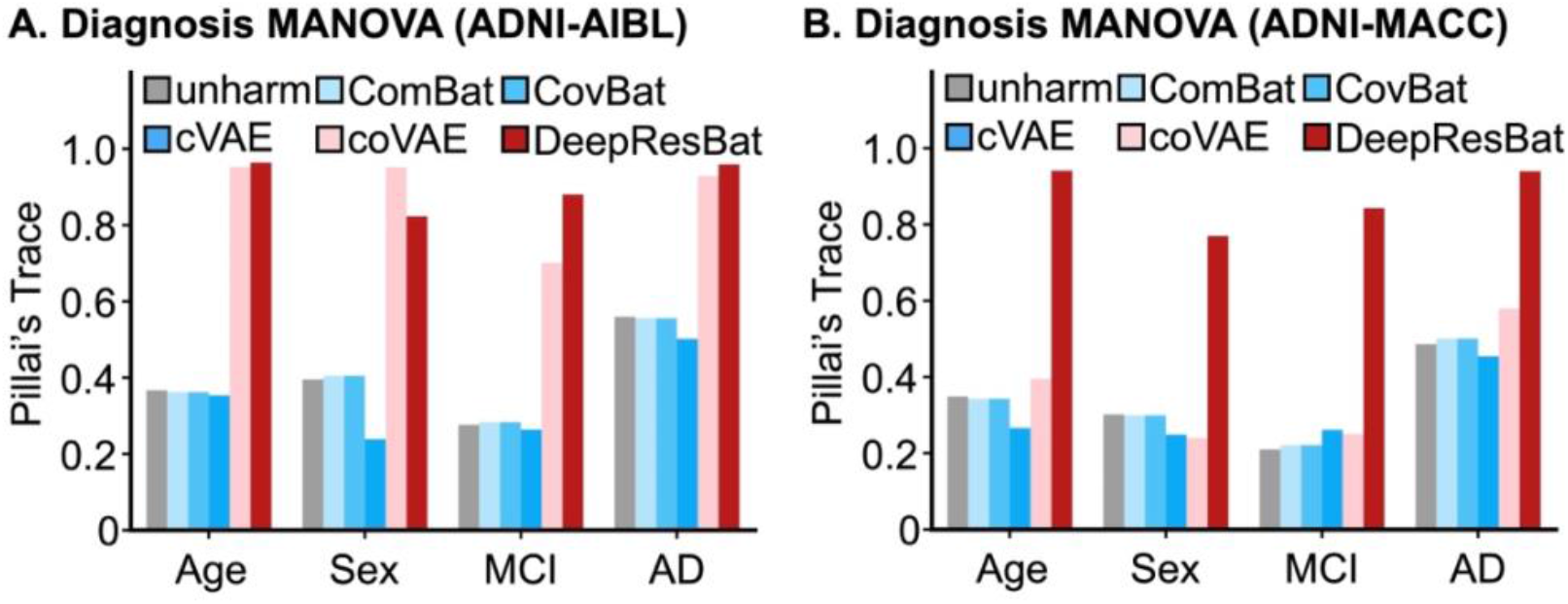
Effect size of MANOVA involving clinical diagnosis. A larger Pillai’s Trace indicates a stronger association, and thus better performance. (A) Bar plot for matched ADNI and AIBL participants. (B) Bar plot for matched ADNI and MACC participants.

In the case of matched ADNI-AIBL participants, DeepResBat yielded stronger brain-covariate associations with respect to no harmonization, ComBat, CovBat and cVAE (Figure 7A). Compared to coVAE, DeepResBat yielded weaker effect sizes for sex, but stronger effect sizes for age, MCI and AD. In the case of matched ADNI-MACC participants, DeepResBat yielded stronger brain-covariate associations with respect to no harmonization and all other baselines (Figure 7B).

Interestingly, coVAE demonstrated inconsistent behavior for association with sex across different cohorts. In the ADNI-AIBL analyses, coVAE harmonized ROIs exhibited a strong association with sex (Figure 7A). By contrast, in the ADNI-MACC analyses, the association with sex was notably weak (Figure 7B). Conversely, DeepResBat displayed more stable performance across cohorts. The inconsistency of coVAE was also present in the univariate GLM analyses (Figures 5 and 6).

Similar conclusions were obtained for the multivariate MANOVA association analyses involving MMSE (Figure S8).

### 3.3 CoVAE, but not DeepResBat, exhibited spurious associations between permuted covariates and harmonized brain volumes

The harmonization models were retrained after permuting all four covariates: age, sex, diagnosis and MMSE (Section 2.10). The GLM association analyses (Section 2.9.1) were then rerun. Figure 8 illustrates the percentage of significant p values (i.e., p < 0.05) across 87 grey matter ROIs from 1000 permutations. More specifically, for each harmonized brain ROI volume, we calculated the percentage of significant p values (i.e., p < 0.05) across 1000 p values corresponding to the 1000 permutations. The expected percentage of significant p values across all grey matter ROIs should be 5%, with a 95% confidence interval (CI) of 3.65% to 6.35% based on the normal approximation of the Binomial 95% CI (Eklund et al., 2016).

**Figure 8.**
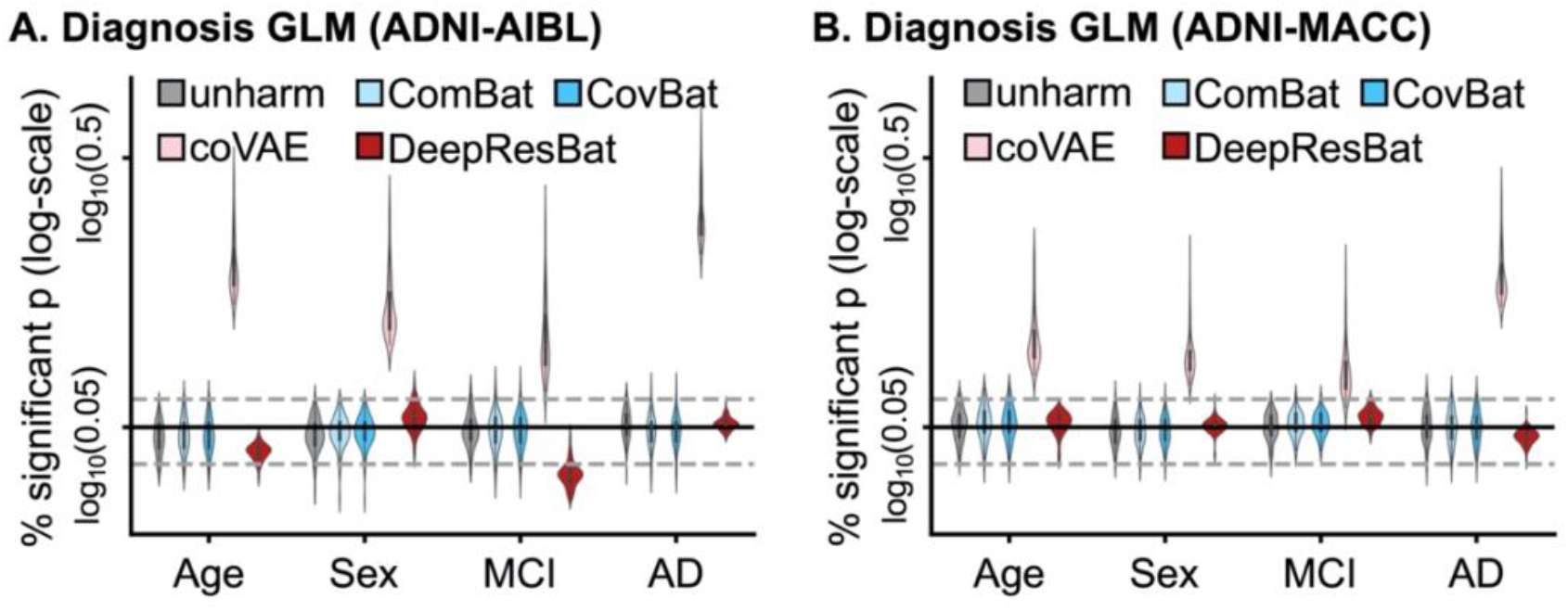
Percentage of significant p values (i.e., p < 0.05) from GLM with clinical diagnosis after 1000 permutations of covariates. More specifically, each data point in the violin plot represents a brain ROI volume. Percentage is calculated based on the number of permutations in which p value of corresponding covariate was significant (i.e., p < 0.05) divided by 1000 permutations. Percentage (vertical axis) is shown on a log scale. The black solid line is the expected percentage (which is 0.05), while the grey dashed lines indicated 95% confidence intervals. (A) GLM analysis involving clinical diagnosis for matched ADNI and AIBL participants. (B) GLM analysis involving clinical diagnosis for matched ADNI and MACC participants.

In both ADNI-AIBL (Figure 8A) and ADNI-MACC (Figure 8B), not performing any harmonization yielded 5% significant p values. ComBat, CovBat and DeepResBat also did not suffer from any spurious associations. However, coVAE yielded an inflated false positive rate in both ADNI-AIBL and ADNI-MACC, since the percentages of significant p values were much greater than 5% for all covariates (Figure 8).

To provide a different visualization of the results, Figure 9 shows the frequency distribution of p values for coVAE and DeepResBat in matched ADNI and AIBL participants. Each line in Figure 9 corresponded to a single brain ROI. The p values were divided into bins with a width of 0.05. Therefore, in the ideal scenario, the distribution of p values should follow a uniform distribution at a height of 0.05. For coVAE (Figure 9A) the frequency of p values within the 0-0.05 bin greatly exceeded 5% for all covariates. For DeepResBat (Figure 9B), there was a uniform distribution of p values for sex and AD. However, the distribution of p values for age and MCI were more conservative with less than 5% of p values in the 0-0.05 bin. Similar results were obtained for matched ADNI and MACC participants (Figure 10). For coVAE (Figure 10A) the frequency of p values within the 0-0.05 bin greatly exceeded 5% for all covariates. For DeepResBat (Figure 10B), the distributions of p values were uniform for all covariates. Visualization of p values distributions for no harmonization, ComBat and CovBat can be found in Figures S9 to S14.

**Figure 9.**
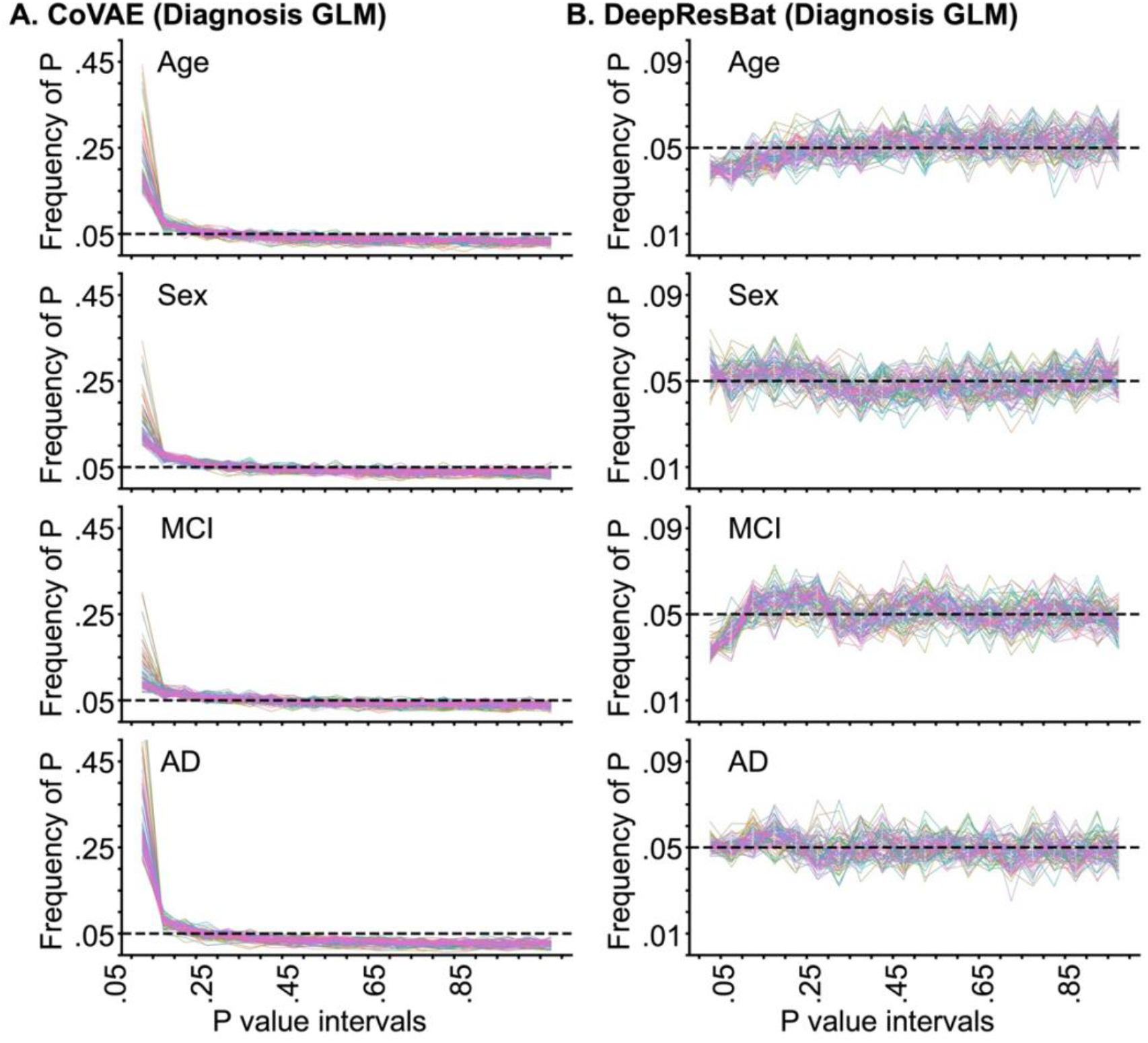
Frequency of p values of coVAE and DeepResBat for matched ADNI and AIBL participants by GLM involving clinical diagnosis based on 1000 permutations. Each line corresponds to a single brain ROI. P values were binned in intervals of 0.05. Therefore, in the ideal scenario, the distributions of p values should follow a uniform distribution with a height of 0.05. (A) Frequency of p values for coVAE. (B) Frequency of p values for DeepResBat.

**Figure 10.**
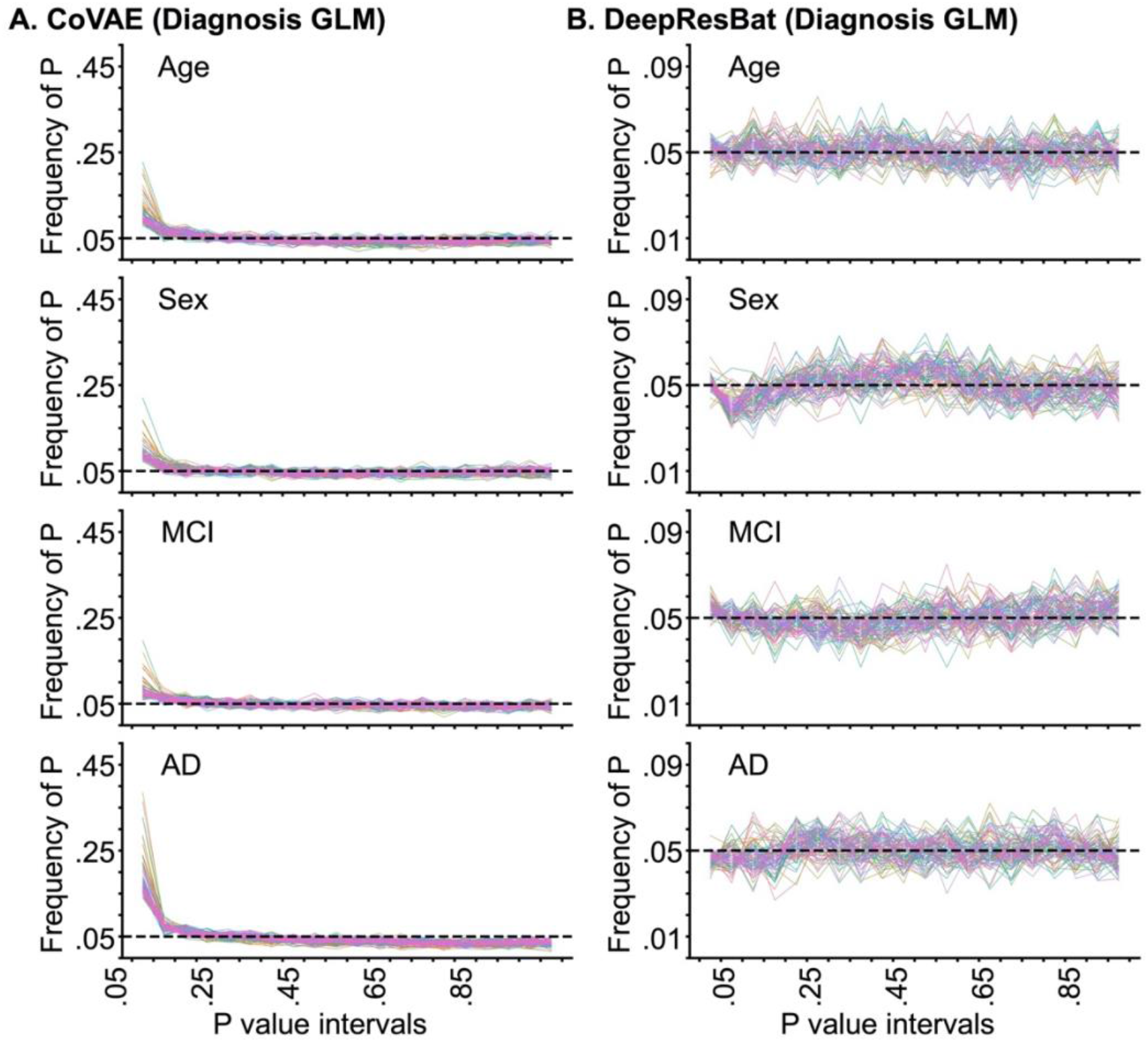
Frequency of p values of coVAE and DeepResBat for matched ADNI and MACC participants by GLM involving clinical diagnosis based on 1000 permutations. Each line corresponds to a single brain ROI. P values were binned in intervals of 0.05. Therefore, in the ideal scenario, the distributions of p values should follow a uniform distribution with a height of 0.05. (A) Frequency of p values for coVAE. (B) Frequency of p values for DeepResBat.

Similar conclusions were obtained for the GLM involving MMSE (Figures S15 to S17). Furthermore, instead of permuting all four covariates, we also considered permuting only clinical diagnosis (Figure S18) or MMSE (Figure S19), yielding similar conclusions.

### 3.4 Further analyses

#### 3.4.1 When harmonizing a pair of datasets, DeepResBat was comparable with ComBat with 50 scans per dataset, and clearly better with 100 scans per dataset

Figure 11 illustrates the dataset prediction accuracies on matched participants harmonized by models fitted with different sample sizes. For both ADNI-AIBL (Figure 11A) and ADNI-MACC (Figure 11B), DeepResBat (red curve) achieves lower dataset prediction accuracies than ComBat (blue curve) from sample size ranging from 50 per dataset to 150 per dataset.

**Figure 11.**
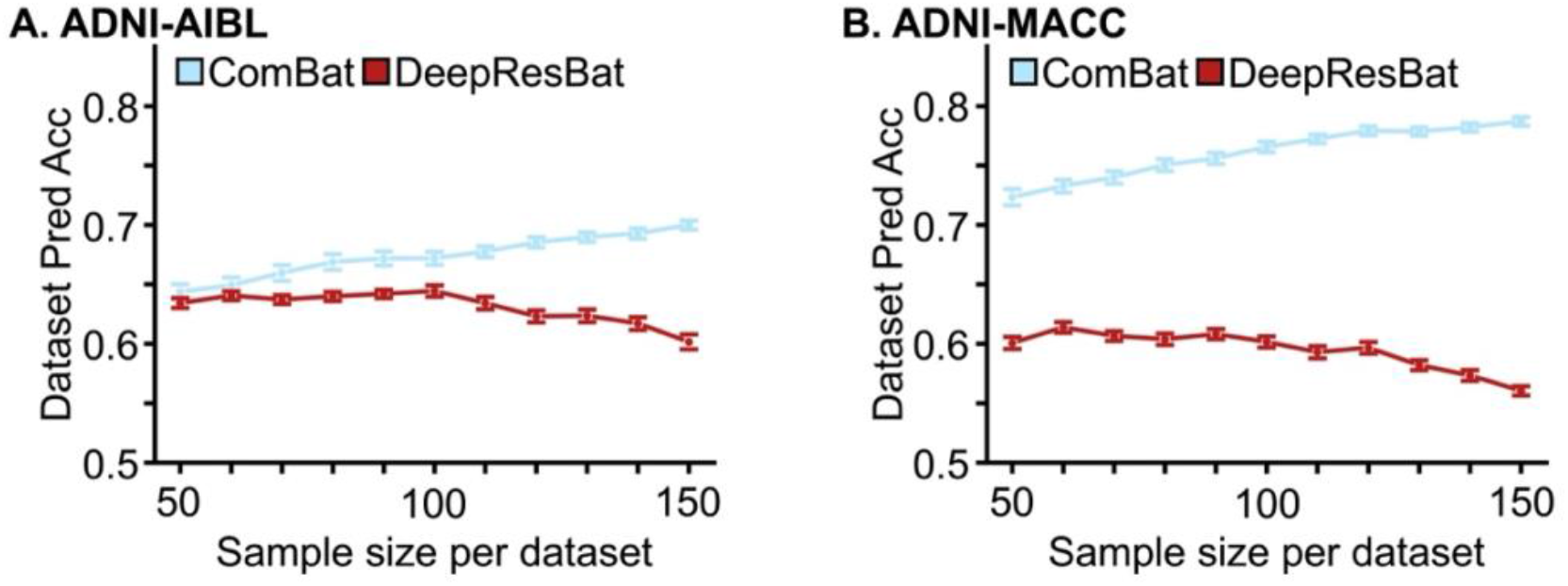
Dataset prediction accuracies for ComBat and DeepResBat with different sample sizes per dataset. Each sampling was repeated 50 times. Error bar represents standard error. (A) Dataset prediction accuracies for matched ADNI and AIBL participants. (B) Dataset prediction accuracies for matched ADNI and MACC participants.

Figures 12 and S20 show the MANOVA effect sizes involving clinical diagnosis and MMSE respectively. In general, ComBat and DeepResBat exhibited similar effect sizes when the sample size per dataset was 50. With 100 MRI scans per dataset, DeepResBat greatly outperformed ComBat and the improvement continued to grow with larger sample sizes.

**Figure 12.**
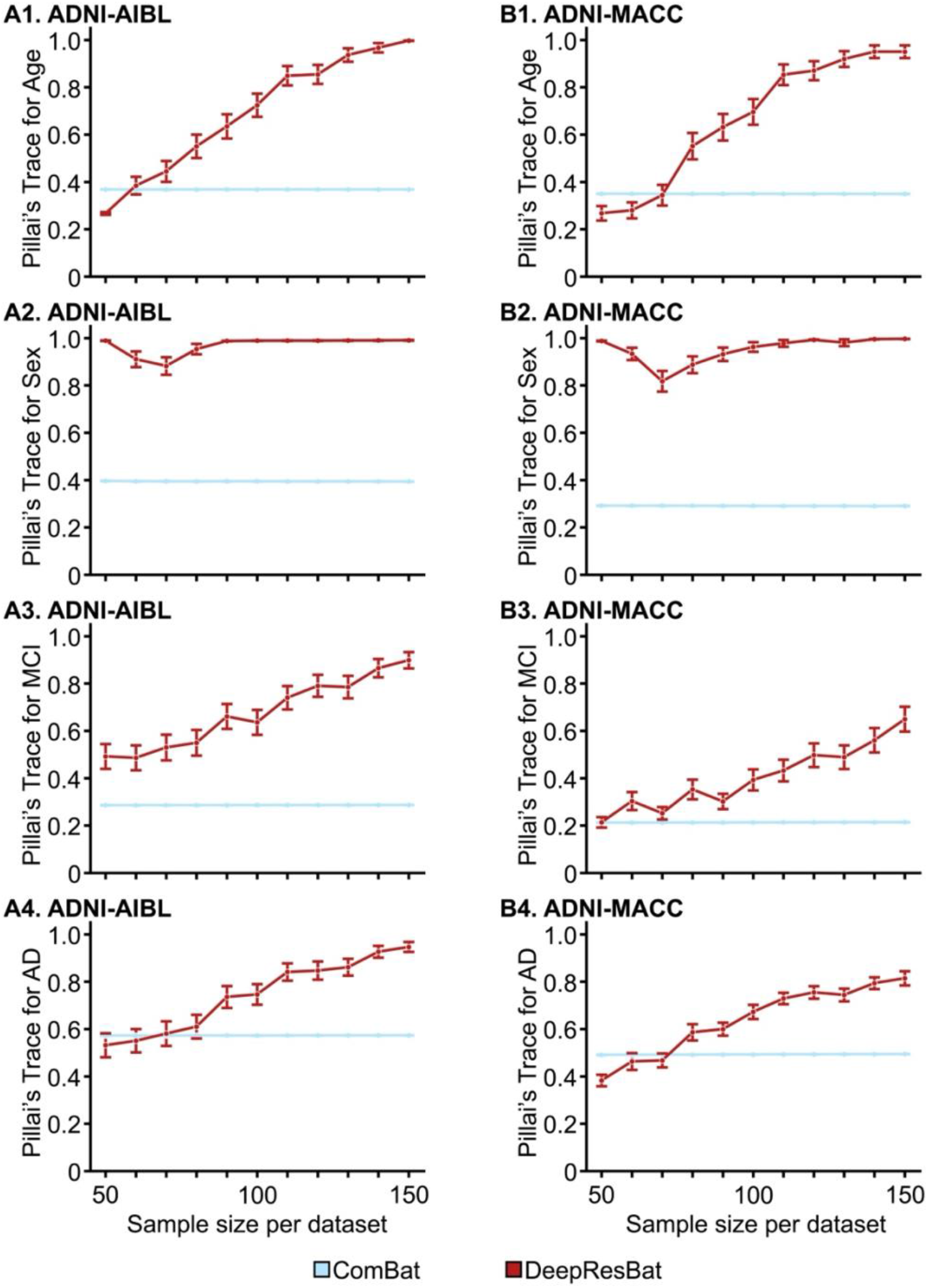
Effect sizes of MANOVA involving clinical diagnosis for ComBat and DeepResBat with different sample size per dataset. Each sampling was repeated 50 times. Error bar corresponds to the standard error. Left column shows the results for harmonizing ADNI and AIBL datasets. Right column shows the results for harmonizing ADNI and MACC datasets. First row corresponds to age associations. Second row corresponds to sex associations. Third row corresponds to MCI associations. Fourth row corresponds to AD diagnosis associations.

#### 3.4.2. DeepResBat outperformed baselines when harmonizing many sites

Figure 13A illustrates the pseudo site prediction accuracies across five folds. Without any harmonization, site prediction accuracy was 0.55, which significantly reduced to 0.29 after ComBat, 0.27 after CovBat and 0.29 after DeepResBat. There was no statistical difference in site prediction accuracies among ComBat, CovBat and DeepResBat.

**Figure 13.**
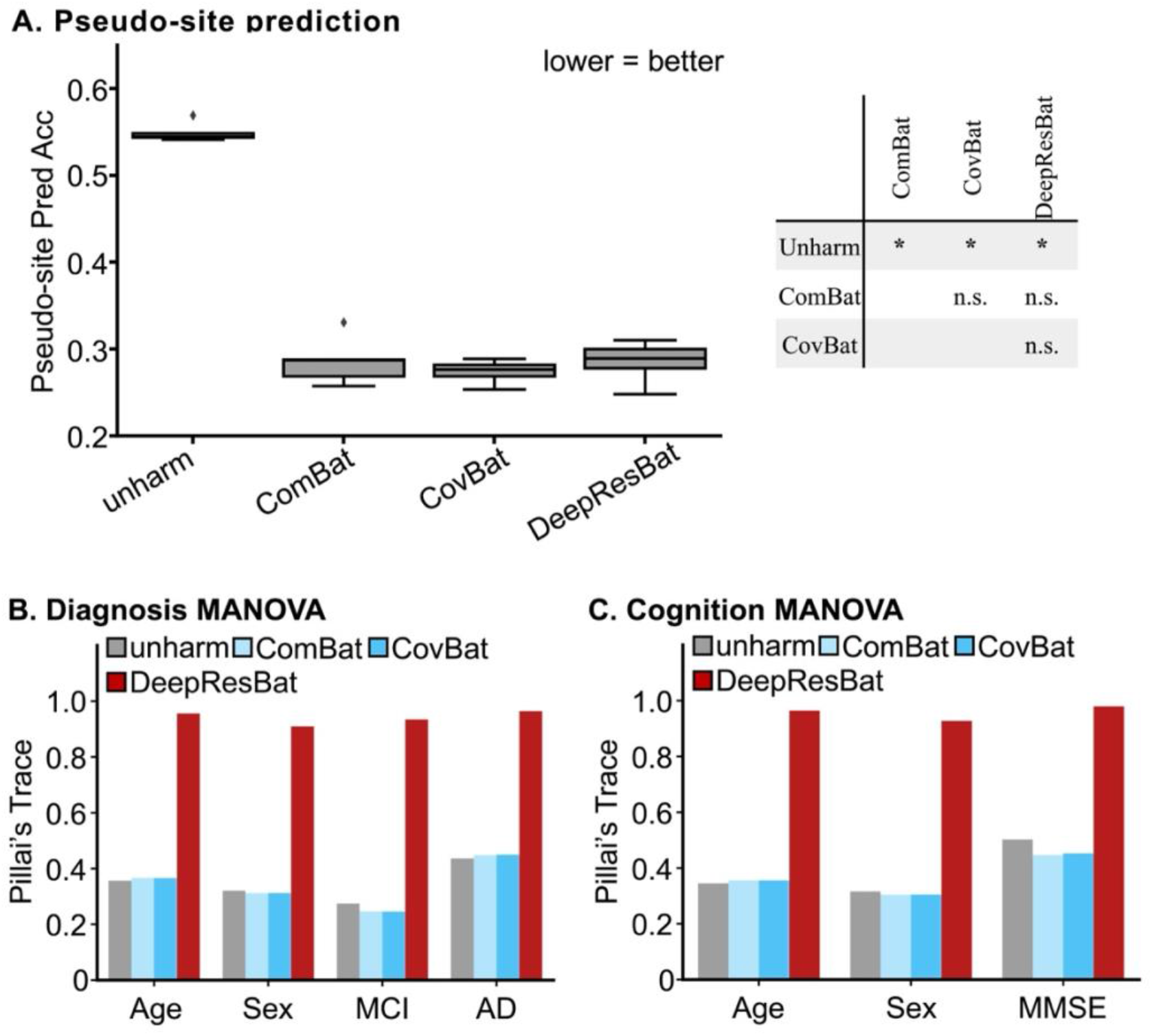
Harmonization of many sites. (A) Dataset prediction accuracies across five folds. Corrected resampled t-test was used as a statistical test to compare different approaches. (B) Effect size of MANOVA involving clinical diagnosis. (C) Effect size of MANOVA involving MMSE. A larger Pillai’s Trace indicates a stronger association, and thus better performance.

Figure 13B and 13C show the multivariate association analysis results by MANOVA involving clinical diagnosis and MMSE respectively. DeepResBat yielded the strongest associations between brain volumes and all covariates. Overall, these results suggest that DeepResBat outperformed ComBat and CovBat when harmonizing many sites.

#### 3.4.3 DeepResBat outperformed baselines on imbalanced test sets

Figure 14 shows the multivariate MANOVA association analysis involving clinical diagnosis in the subsampled test sets with highly imbalanced covariates. DeepResBat exhibited stronger associations between covariates and brain volumes than no harmonization, ComBat and CovBat for all scenarios. Similar conclusions were obtained for the multivariate MANOVA association analyses involving MMSE (Figure S21).

**Figure 14.**
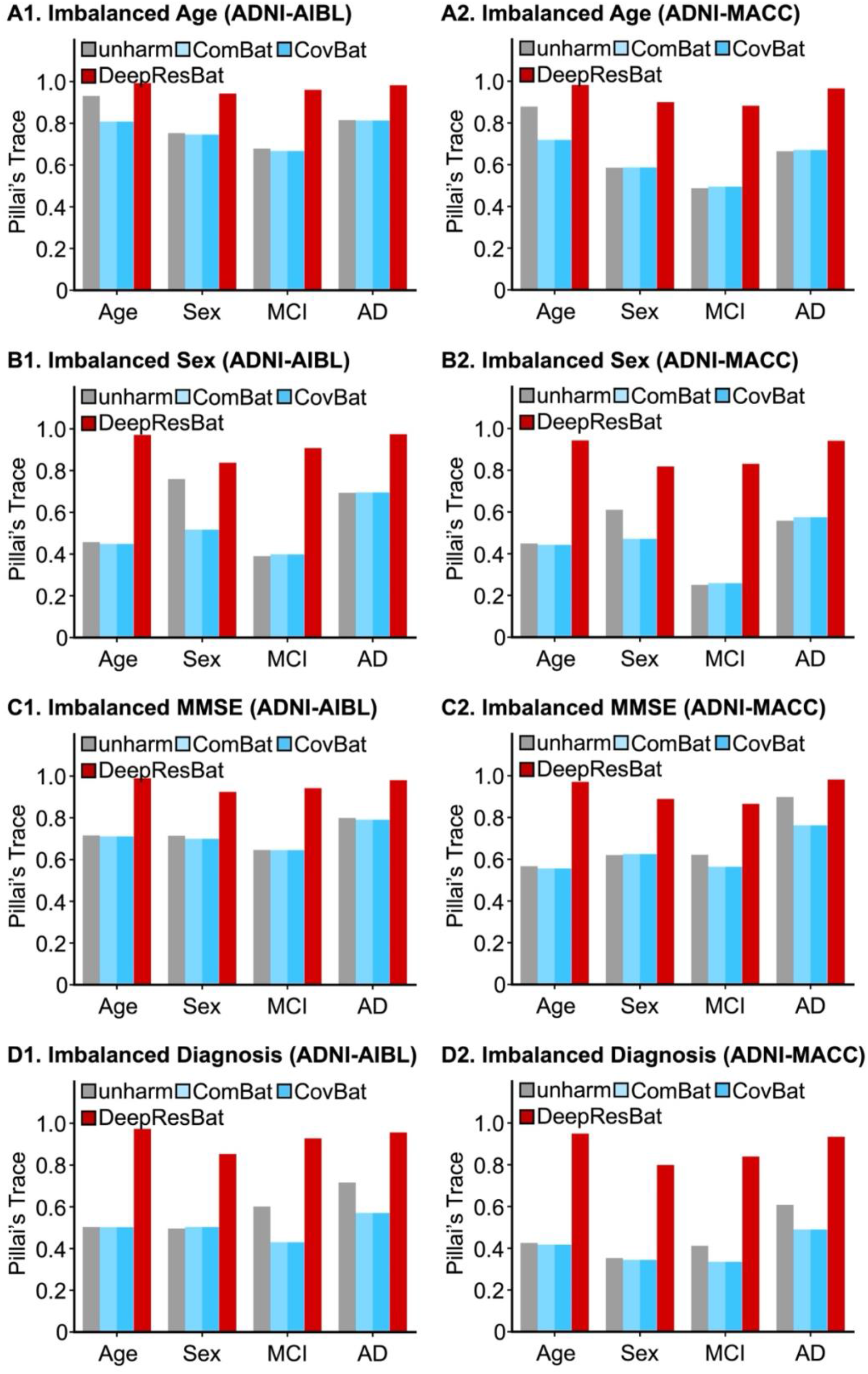
Effect size of MANOVA involving clinical diagnosis in test sets involving highly imbalanced covariate distributions. A larger Pillai’s Trace indicates a stronger association, and thus better performance. The left column corresponding to harmonizing ADNI and AIBL. The right column corresponding to harmonizing ADNI and MACC. (A1) MANOVA effect sizes for ADNI-AIBL test set with highly imbalanced age distributions. (A2) Same as A1 but for ADNI and MACC. (B1) MANOVA effect sizes for ADNI-AIBL test set with highly imbalanced sex distributions. (B2) Same as B1 but for ADNI and MACC. (C1) MANOVA effect sizes for ADNI-AIBL test set with highly imbalanced MMSE distributions. (C2) Same as C1 but for ADNI and MACC. (D1) MANOVA effect sizes for ADNI-AIBL test set with highly imbalanced diagnosis distributions. (D2) Same as B1 but for ADNI and MACC.

## 4 Discussion

Current deep learning approaches for harmonization do not explicitly account for covariate distribution differences across datasets. As discussed in Section 2.1, ignoring covariates can lead to theoretically worse harmonization outcomes. We then proposed two DNN-based harmonization approaches, coVAE and DeepResBat, which explicitly accounted for covariate distribution differences across datasets. We demonstrated that DeepResBat outperformed mixed effects and deep learning baselines across three evaluation experiments involving three large-scale MRI datasets.

More specifically, without any harmonization, XGBoost was able to predict almost perfectly which dataset a participant’s MRI volumes came from (Figure 4). After harmonization with mixed effects models (ComBat and CovBat), dataset classification accuracies dropped significantly, suggesting that ComBat and CovBat were able to remove some dataset differences. DNN-based harmonization approaches further reduced the classification accuracies, suggesting even greater removal of dataset differences.

However, the removed dataset differences might contain important biological information, which should not be removed. Therefore, in the second experiment, we evaluated the strength of associations between the harmonized brain volumes and covariates (age, sex, MMSE and clinical diagnosis). Across both univariate GLM and multivariate MANOVA (Figures 5 to 7), we found that coVAE and DeepResBat yielded stronger associations between brain volumes and covariates. This suggests that coVAE and DeepResBat were retaining important biological information while removing undesirable dataset differences. Interestingly, DeepResBat was also more sensitive than coVAE, except for the association between brain volumes and sex in the matched ADNI and AIBL participants. Finally, cVAE exhibited weaker associations than ComBat, CovBat and even no harmonization (Figure 7), suggesting that cVAE was removing significant biological information in addition to unwanted dataset differences, thus providing empirical support for the theoretical discussion in Section 2.1.

Given the flexible nature of DNNs, we were concerned that explicitly accounting for covariates could lead to spurious associations between harmonized brain volumes and covariates when no association existed. Our permutation test (Figures 8 to 10) supported our concerns in the case of coVAE. Although coVAE provided a natural (and in our opinion, elegant) extension of cVAE, we found significant false positive rates for coVAE. On the other hand, DeepResBat was able to exhibit an expected amount of false positives, consistent with less flexible mixed effects models (ComBat and CovBat).

Together, the three evaluation experiments suggest that DeepResBat is an effective deep learning alternative to ComBat. DeepResBat consisted of three steps: (1) regressed out the effects of covariates from the brain volumes, (2) followed by harmonizing the residuals, (3) and then adding the effects of covariates back to the harmonized residuals. Future research could investigate whether these three steps can be combined into a single optimization procedure by minimizing fitting Eq. (10) directly. However, we note that fitting Eq. (10) directly might lead to overfitting, yielding false positive issues, similar to coVAE. While our current implementation of DeepResBat utilized XGBoost to estimate covariate effects, other nonlinear regression approaches can be used. Furthermore, instead of using cVAE in the harmonization step, cVAE can be replaced with other harmonization approaches, such as generative adversarial networks (Bashyam et al., 2021).

We performed three additional analyses to address the practical considerations of using DeepResBat. When harmonizing a pair of datasets, we found that DeepResBat and ComBat had similar performance when there were 50 T1 scans per dataset, while DeepResBat was much better than ComBat when there were at least 90 to 100 T1 scans per dataset (Figures 11, 12 and S20). This suggests that for simplicity, ComBat can be used when sample size per dataset is 50 scans or less, while DeepResBat should definitely be used when sample size reaches 100 scans per dataset.

One advantage of cVAE is that the approach easily works for more than two datasets by extending the one-hot encoding of sites. Our analysis harmonizing 21 pseudo sites with a wide range of sample size and covariate distributions (Figure 3) suggests that DeepResBat greatly outperformed ComBat and CovBat in terms of strengthening the associations between covariates and brain volumes, while removing site differences (Figure 13). This suggests the utility of DeepResBat for mega-analysis involving posthoc data collation across many sites, such as ENIGMA (Larivière et al., 2022; van Velzen et al., 2020), Global Brain Consortium (Valdes-Sosa et al., 2021, 2022) and Lifespan Brain Chart Consortium (Bethlehem et al., 2022) as well as large-scale projects involving coordinated data collection across many sites, such as ABCD (Volkow et al., 2018) and Human Connectome Project Lifespan (Van Essen et al., 2012; Harms et al., 2018; Somerville et al., 2018) studies.

Finally, DeepResBat can be trained on datasets with somewhat unbalanced covariate distributions, which can then be applied to test data with highly imbalanced covariate distributions (Figures 14 and S21). Given the relatively small sample size requirements (Figures 11, 12 and S20), a user could train on a relatively small subset of data, and then applied the resulting DeepResBat model to the remaining data. Alternatively, users can also train and apply DeepResBat on the full set of data.

The usage of DeepResBat is relatively straightforward. Examples have been provided in our GitHub repository to guide users for applying DeepResBat. Users just need to specify the data matrix (# scans × # features) to harmonize, as well as the site and covariates for each scan. Our wrapper function will automatically perform the hyperparameter search for both XGBoost and cVAE. For an 8000 × 108 data matrix, training requires around 4GB of GPU memory and 20 hours on a RTX3090 (inclusive of hyperparameter search). Alternatively, users have the option to only perform hyperparameter search for XGBoost and use the default cVAE hyperparameters that we have provided. In our experience, cVAE is relatively robust to hyperparameter choices. In this scenario, for an 8000 × 108 data matrix, training requires around 1GB of GPU memory and 10 min on a RTX3090 (inclusive of XGBoost hyperparameter search). Inference only requires a forward pass through the DNN and applying the XGBoost models, so it is extremely fast.

A drawback of DeepResBat is that our current implementation operates on summary measures (e.g., volumes or thickness), rather than at the image level (Zuo et al., 2021; Cackowski et al., 2023). Therefore, the harmonization procedure needs to be repeated for different summary measures (e.g., using a different brain parcellation). However, this disadvantage also means that DeepResBat can be applied to harmonize not just T1 MRI data, but also any tabular data. DeepResBat might therefore be applicable to imaging modalities, such as diffusion MRI or PET, where summary measures for certain brain regions might be computed for further analyses.

However, one challenge is that the runtime of the covariate effects estimation (Step 1 of DeepResBat) will increase linearly as the number of input variables increases. Similarly, the runtime of the multilayer perceptron utilized in cVAE (Step 2 of DeepResBat) will increase linearly as the number of input variables increases. Overall, the runtime barrier would be challenging for harmonizing large number of variables, e.g., the tens of thousands of variables in a typical functional connectivity matrix (Yan et al., 2023; P. Chen et al., 2024) In this scenario, instead of a multilayer perceptron implemented in the current study, a more efficient neural network architecture that exploits the natural structure of the data can be considered.

Beyond imaging data, DeepResBat can be broadly applicable to any field where instrumental harmonization is necessary, e.g., when combining data across sensors from different satellites (Asim et al., 2023) and radiosounding stations (Madonna et al., 2022). To provide a more detailed example, the ComBat model was originally proposed for harmonizing batches of microarray experiments (Johnson et al., 2007). A biologist might be interested in comparing the gene expression profiles of diseased and healthy tissues. Due to throughput limitations, these tissue samples might be processed in batches days or months apart, resulting in non-biological batch effects. Batch effects might also arise when combining data across different labs, array types, or platforms (Rhodes et al., 2004). Therefore, such batch effects need to be removed via harmonization. When performing harmonization, the experimental conditions (e.g., disease status of the tissue) are relevant covariates, whose effects should be retained after harmonization. In other words, DeepResBat should also be applicable to microarray data.

## 5 Conclusion

In this study, we demonstrate the importance of incorporating covariates during harmonization. We propose two deep learning models, coVAE and DeepResBat, that account for covariate distribution differences across datasets. coVAE extends cVAE by concatenating covariates and site information with latent representations, while DeepResBat adopts a residual framework inspired by the classical ComBat framework. We found that coVAE introduces spurious associations between anatomical MRI and unrelated covariates, while DeepResBat effectively mitigates this false positive issue. Furthermore, DeepResBat outperformed ComBat, CovBat and cVAE in terms of removing dataset differences, while retaining biological effects of interest.

## Acknowledgment

Our research is currently supported by the NUS Yong Loo Lin School of Medicine (NUHSRO/2020/124/TMR/LOA), the Singapore National Medical Research Council (NMRC) LCG (OFLCG19May-0035), NMRC CTG-IIT (CTGIIT23jan-0001), NMRC OF-IRG (OFIRG24jan-0030), NMRC STaR (STaR20nov-0003), Singapore Ministry of Health (MOH) Centre Grant (CG21APR1009), the Temasek Foundation (TF2223-IMH-01), and the United States National Institutes of Health (R01MH120080 & R01MH133334). Our computational work was partially performed on resources of the National Supercomputing Centre, Singapore (https://www.nscc.sg). Any opinions, findings and conclusions or recommendations expressed in this material are those of the authors and do not reflect the views of the funders. Data collection and sharing for this project was funded by the Alzheimer’s Disease Neuroimaging Initiative (ADNI) (National Institutes of Health Grant U01 AG024904) and DOD ADNI (Department of Defense award number W81XWH-12-2-0012). ADNI is funded by the National Institute on Aging, the National Institute of Biomedical Imaging and Bioengineering, and through generous contributions from the following: AbbVie, Alzheimer’s Association; Alzheimer’s Drug Discovery Foundation; Araclon Biotech; BioClinica, Inc.; Biogen; Bristol-Myers Squibb Company; CereSpir, Inc.; Cogstate; Eisai Inc.; Elan Pharmaceuticals, Inc.; Eli Lilly and Company; EuroImmun; F. Hoffmann-La Roche Ltd and its affiliated company Genentech, Inc.; Fujirebio; GE Healthcare; IXICO Ltd.;Janssen Alzheimer Immunotherapy Research & Development, LLC.; Johnson & Johnson Pharmaceutical Research & Development LLC.; Lumosity; Lundbeck; Merck & Co., Inc.;Meso Scale Diagnostics, LLC.; NeuroRx Research; Neurotrack Technologies; Novartis Pharmaceuticals Corporation; Pfizer Inc.; Piramal Imaging; Servier; Takeda Pharmaceutical Company; and Transition Therapeutics. The Canadian Institutes of Health Research is providing funds to support ADNI clinical sites in Canada. Private sector contributions are facilitated by the Foundation for the National Institutes of Health (www.fnih.org). The grantee organization is the Northern California Institute for Research and Education, and the study is coordinated by the Alzheimer’s Therapeutic Research Institute at the University of Southern California. ADNI data are disseminated by the Laboratory for Neuro Imaging at the University of Southern California. The AIBL study would like to thank all of the participants who took part in the study and the clinicians who referred participants. The AIBL study (www.AIBL.org.au) is a consortium between Austin Health, CSIRO, Edith Cowan University, the Florey Institute (The University of Melbourne), and the National Ageing Research Institute. Partial financial support provided by the Alzheimer’s Association (US), the Alzheimer’s Drug Discovery Foundation, an Anonymous foundation, the Science and Industry Endowment Fund, the Dementia Collaborative Research Centres, the Victorian Government’s Operational Infrastructure Support program, the McCusker Alzheimer’s Research Foundation, the National Health and Medical Research Council, and the Yulgilbar Foundation. Numerous commercial interactions have supported data collection and analysis. In-kind support has also been provided by Sir Charles Gairdner Hospital, CogState Ltd., Hollywood Private Hospital, the University of Melbourne, and St Vincent’s Hospital

## CRediT authorship contribution statement

**Lijun An**: Conceptualization, Data curation, Formal analysis, Investigation, Methodology, Project administration, Software, Visualization, Writing – original draft, Writing – review & editing. **Chen Zhang**: Software, Validation, Visualization, Writing – review & editing. **Naren Wulan**: Investigation, Software, Validation, Visualization, Writing – review & editing. **Shaoshi Zhang**: Investigation, Software, Validation, Visualization, Writing – review & editing. **Pansheng Chen**: Software, Validation, Visualization, Writing – review & editing. **Fang Ji**: Resource, Writing – review & editing. **Kwun Kei Ng**: Investigation, Writing – review & editing. **Christopher Chen**: Resource, Writing – review & editing. **Juan Helen Zhou**: Resource, Writing – review & editing. **B.T. Thomas Yeo**: Conceptualization, Formal analysis, Funding acquisition, Investigation, Methodology, Resource, Supervision, Visualization, Writing – original draft, Writing – review & editing.

