## Supplementary for "DeepResBat: deep residual batch harmonization accounting for covariate distribution differences"

### Supplementary Material

| Vendor | Scanner Model | Field Strength | Number of scans |
|---|---|---|---|
| GE | Discovery | 3T | 595 |
| | Genesis Signa | 3T | 273 |
| | Signa Excite | 1.5T | 838 |
| | | 3T | 30 |
| | Signa HDx | 1.5T | 464 |
| | | 3T | 42 |
| | Signa HDxt | 1.5T | 212 |
| | | 3T | 405 |
| Philips | Achieva | 1.5T | 67 |
| | | 3T | 481 |
| | Gemini | 3T | 32 |
| | Gyroscan Intera | 1.5T | 12 |
| | Gyroscan NT | 1.5T | 2 |
| | Ingenia | 3T | 84 |
| | Ingenuity | 3T | 18 |
| | Intera | 1.5T | 319 |
| | | 3T | 216 |
| | Intera Achieva | 1.5T | 6 |
| | | 3T | 1 |
| Siemens | Allegra | 3T | 48 |
| | Avanto | 1.5T | 385 |
| | Biograph | 3T | 12 |
| | Espree | 1.5T | 22 |
| | NUMARIS/4 | 1.5T | 2 |
| | Prisma | 3T | 2 |
| | Prisma_fit | 3T | 3 |
| | Skyra | 3T | 274 |
| | Sonata | 1.5T | 371 |
| | SonataVision | 1.5T | 25 |
| | Symphony | 1.5T | 547 |
| | SymphonyTim | 1.5T | 88 |
| | Trio | 3T | 107 |
| | TrioTim | 3T | 1371 |
| | Verio | 3T | 601 |

**Table S1. Scanner information for 7955 scans in ADNI dataset.**

| Vendor | Scanner Model | Field Strength | Number of scans |
|---|---|---|---|
| Siemens | Avanto | 1.5T | 241 |
| | TrioTim | 3T | 558 |
| | Verio | 3T | 134 |

**Table S2. Scanner information for 933 scans in AIBL dataset.**

| Dataset | Min # scans | Max # scans | Mean # scans | Median # scans |
|---|---|---|---|---|
| ADNI | 1 | 11 | 4.59 | 5 |
| AIBL | 1 | 4 | 1.88 | 1 |
| MACC | 1 | 3 | 2.15 | 2 |

**Table S3. Summary statistics of the number of MRI scans per participant.**

| | Timepoint | ADNI value | AIBL value | P value |
|---|---|---|---|---|
| **AGE** | 1 | 71.0±5.5 | 70.8±5.3 | 0.96 |
| | 2 | 72.5±5.5 | 72.6±5.5 | 0.98 |
| | 3 | 74.2±5.5 | 73.9±5.6 | 0.93 |
| | 4 | 75.7±5.5 | 75.6±5.5 | 0.99 |
| **MMSE** | 1 | 29.3±0.9 | 29.2±0.9 | 1.00 |
| | 2 | 29.5±0.5 | 29.5±0.5 | 1.00 |
| | 3 | 29.7±0.5 | 29.7±0.5 | 1.00 |
| | 4 | 29.5±0.8 | 29.5±0.8 | 1.00 |
| **AD diagnosis** | 1 | 100%-0%-0% | 100%-0%-0% | 1.00 |
| | 2 | 100%-0%-0% | 100%-0%-0% | 1.00 |
| | 3 | 100%-0%-0% | 100%-0%-0% | 1.00 |
| | 4 | 100%-0%-0% | 100%-0%-0% | 1.00 |
| **Sex** | - | 50% | 50% | 1.00 |

**Table S4.** ADNI-AIBL matching results for participants having 4 time points (scans). For clinical diagnosis in the table, the percentage is showed as CN%-MCI%-AD%. For sex in the table, the portion is the ratio of male subjects. For age, the p value was calculated with a two-sample t-test. For MMSE, the p value was calculated from a Kolmogorov–Smirnov test. For sex/AD diagnosis, the p value was calculated from the chi-square goodness of fit test.

| | Timepoint | ADNI value | AIBL value | P value |
|---|---|---|---|---|
| AGE | 1 | 73.3±3.3 | 73.1±3.3 | 0.96 |
| | 2 | 74.8±3.3 | 75.2±3.3 | 0.94 |
| | 3 | 76.3±3.3 | 76.1±3.3 | 0.97 |
| MMSE | 1 | 29.0±0.0 | 20.0±0.0 | 1.00 |
| | 2 | 30.0±0.0 | 30.0±0.0 | 1.00 |
| | 3 | 30.0±0.0 | 30.0±0.0 | 1.00 |
| AD diagnosis | 1 | 100%-0%-0% | 100%-0%-0% | 1.00 |
| | 2 | 100%-0%-0% | 100%-0%-0% | 1.00 |
| | 3 | 100%-0%-0% | 100%-0%-0% | 1.00 |
| Sex | - | 50% | 50% | 1.00 |

**Table S5.** ADNI-AIBL matching results for participants having 3 time points (scans). For clinical diagnosis in the table, the percentage is showed as CN%-MCI%-AD%. For sex in the table, the portion is the ratio of male subjects. For age, the p value was calculated with a two-sample t-test. For MMSE, the p value was calculated from a Kolmogorov–Smirnov test. For sex/AD diagnosis, the p value was calculated from the chi-square goodness of fit test.

| | Timepoint | ADNI value | AIBL value | P value |
|---|---|---|---|---|
| AGE | 1 | 74.4±9.8 | 74.5±9.8 | 0.99 |
| | 2 | 76.1±9.8 | 76.1±9.9 | 0.99 |
| MMSE | 1 | 27.9±2.8 | 27.9±2.8 | 1.00 |
| | 2 | 27.8±2.8 | 27.8±2.8 | 1.00 |
| AD diagnosis | 1 | 57%-43%-0% | 57%-43%-0% | 1.00 |
| | 2 | 57%-43%-0% | 57%-43%-0% | 1.00 |
| Sex | - | 88% | 88% | 1.00 |

**Table S6.** ADNI-AIBL matching results for participants having 2 time points (scans). For clinical diagnosis in the table, the percentage is showed as CN%-MCI%-AD%. For sex in the table, the portion is the ratio of male subjects. For age, the p value was calculated with a two-sample t-test. For MMSE, the p value was calculated from a Kolmogorov–Smirnov test. For sex/AD diagnosis, the p value was calculated from the chi-square goodness of fit test.

| | Timepoint | ADNI value | AIBL value | P value |
|---|---|---|---|---|
| **AGE** | 1 | 74.8±5.9 | 74.8±5.9 | 1.00 |
| **MMSE** | 1 | 27.3±3.9 | 27.3±3.9 | 0.98 |
| **AD diagnosis** | 1 | 68%-19%-13% | 68%-19%-13% | 1.00 |
| **Sex** | - | 43% | 43% | 1.00 |

**Table S7.** ADNI-AIBL matching results for participants having 1 time point (scan). For clinical diagnosis in the table, the percentage is showed as CN%-MCI%-AD%. For sex in the table, the portion is the ratio of male subjects. For age, the p value was calculated with a two-sample t-test. For MMSE, the p value was calculated from a Kolmogorov–Smirnov test. For sex/AD diagnosis, the p value was calculated from the chi-square goodness of fit test.

| | Timepoint | ADNI value | MACC value | P value |
|---|---|---|---|---|
| **AGE** | 1 | 71.5±6.8 | 72.3±6.7 | 0.67 |
| | 2 | 73.5±6.8 | 73.8±6.8 | 0.91 |
| | 3 | 75.9±6.9 | 75.5±6.6 | 0.81 |
| **MMSE** | 1 | 26.9±3.7 | 27.0±3.5 | 1 |
| | 2 | 26.1±4.5 | 26.1±4.5 | 1 |
| | 3 | 24.9±6.3 | 25.2±6.3 | 0.99 |
| **AD diagnosis** | 1 | 39%-46%-15% | 36%-54%-10% | 0.72 |
| | 2 | 43%-36%-21% | 46%-36%-18% | 0.88 |
| | 3 | 43%-36%-21% | 46%-32%-22% | 0.91 |
| **Sex** | - | 57% | 57% | 1.00 |

**Table S8.** ADNI-MACC matching results for participants having 3 time points (scans). For clinical diagnosis in the table, the percentage is showed as CN%-MCI%-AD%. For sex in the table, the portion is the ratio of male subjects. For age, the p value was calculated with a two-sample t-test. For MMSE, the p value was calculated from a Kolmogorov–Smirnov test. For sex/AD diagnosis, the p value was calculated from the chi-square goodness of fit test.

| | Timepoint | ADNI value | MACC value | P value |
|---|---|---|---|---|
| **AGE** | 1 | 73.6±5.7 | 73.9±5.6 | 0.78 |
| | 2 | 75.8±5.6 | 75.5±5.6 | 0.71 |
| **MMSE** | 1 | 24.7±4.9 | 24.8±4.6 | 1 |
| | 2 | 23.4±6.9 | 23.5±6.6 | 1 |
| **AD diagnosis** | 1 | 35%-38%-27% | 35%-40%-25% | 0.80 |
| | 2 | 37%-30%-33% | 37%-35%-28% | 0.49 |
| **Sex** | - | 51% | 58% | 0.20 |

**Table S9.** ADNI-MACC matching results for participants having 2 time points (scans). For clinical diagnosis in the table, the percentage is showed as CN%-MCI%-AD%. For sex in the table, the portion is the ratio of male subjects. For age, the p value was calculated with a two-sample t-test. For MMSE, the p value was calculated from a Kolmogorov–Smirnov test. For sex/AD diagnosis, the p value was calculated from the chi-square goodness of fit test

| | Timepoint | ADNI value | MACC value | P value |
|---|---|---|---|---|
| **AGE** | 1 | 75.7±6.7 | 75.7±6.7 | 0.97 |
| **MMSE** | 1 | 21.0±5.9 | 21.0±5.9 | 1 |
| **AD diagnosis** | 1 | 14%-34%-52% | 14%-38%-48% | 0.64 |
| **Sex** | - | 52% | 56% | 0.34 |

**Table S10.** ADNI-MACC matching results for participants having 1 time points (scans). For clinical diagnosis in the table, the percentage is showed as CN%-MCI%-AD%. For sex in the table, the portion is the ratio of male subjects. For age, the p value was calculated with a two-sample t-test. For MMSE, the p value was calculated from a Kolmogorov–Smirnov test. For sex/AD diagnosis, the p value was calculated from the chi-square goodness of fit test.

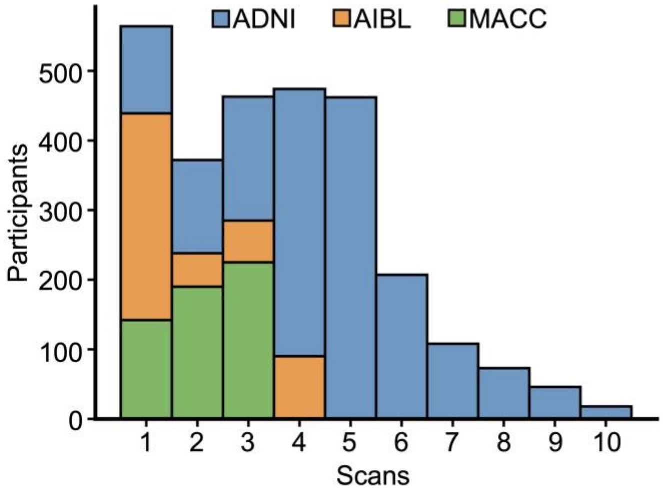


**Figure S1. Distribution of the number of MRI scans per participant.**

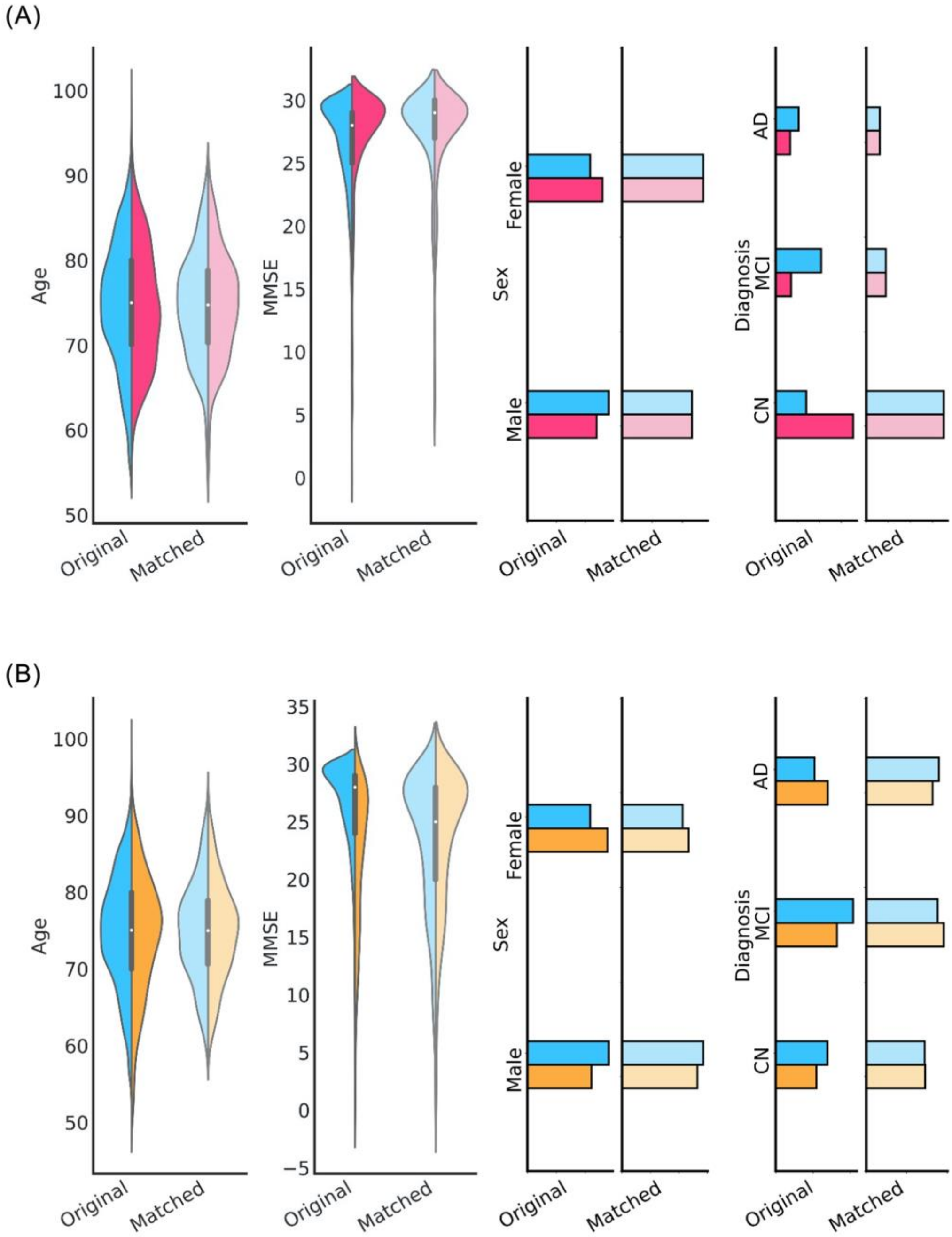


**Figure S2. Age, MMSE, sex and clinical diagnosis distributions before and after matching.** (A) Distributions of age, sex, MMSE and clinical diagnosis for ADNI (blue) and AIBL (red). Differences in the attributes between ADNI and AIBL were not significant after matching. (B) Distributions of age, sex, MMSE and clinical diagnosis for ADNI (blue) and MACC (yellow). Differences in the attributes between ADNI and MACC were not significant after matching. P values showing the quality of the matching procedure are found in Tables S3 to S9.

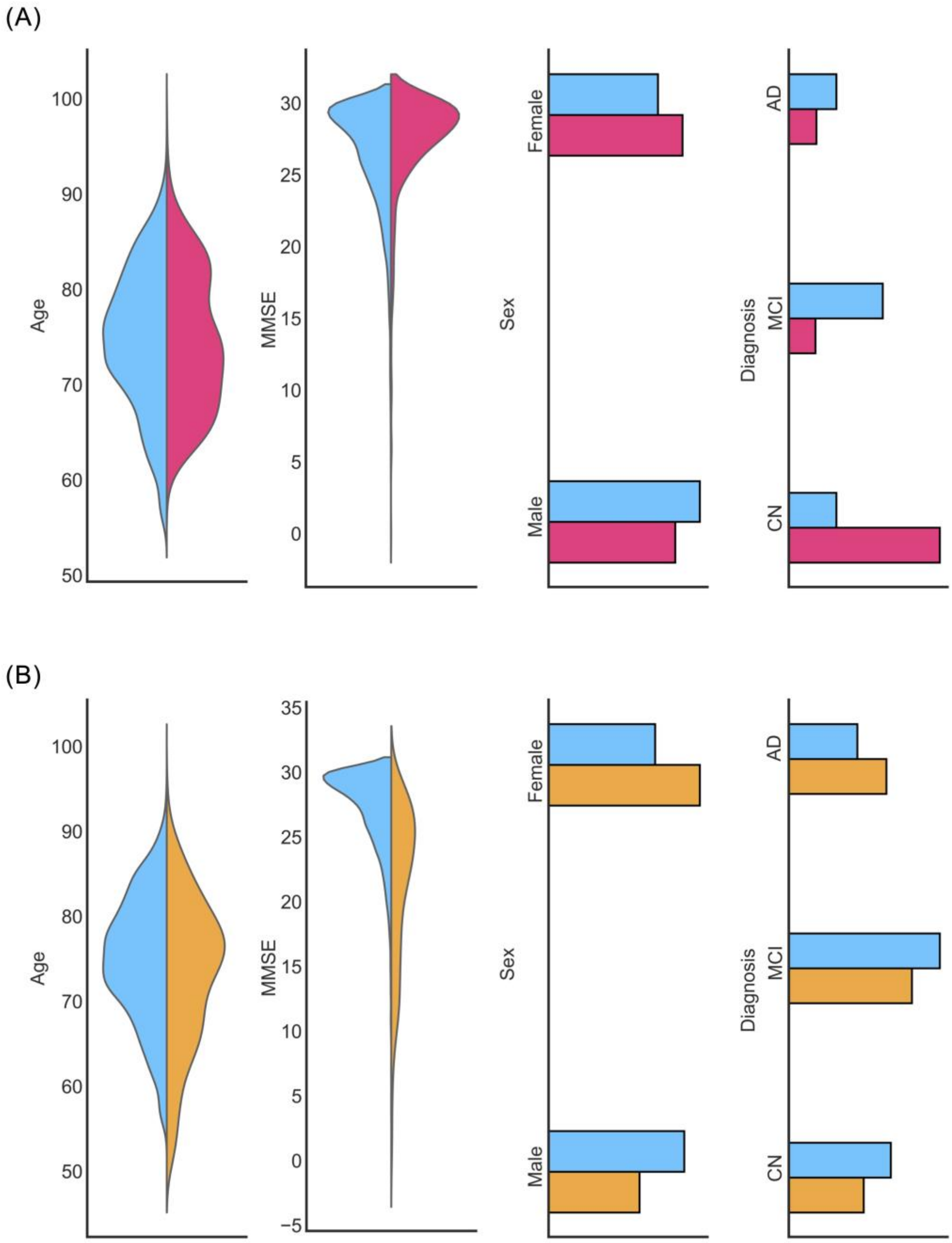


**Figure S3. Age, MMSE, sex and clinical diagnosis distributions of unmatched participants.** (A) Distributions of age, sex, MMSE and clinical diagnosis for unmatched ADNI (blue) and unmatched AIBL (red). (B) Distributions of age, sex, MMSE and clinical diagnosis for unmatched ADNI (blue) and unmatched MACC (yellow).

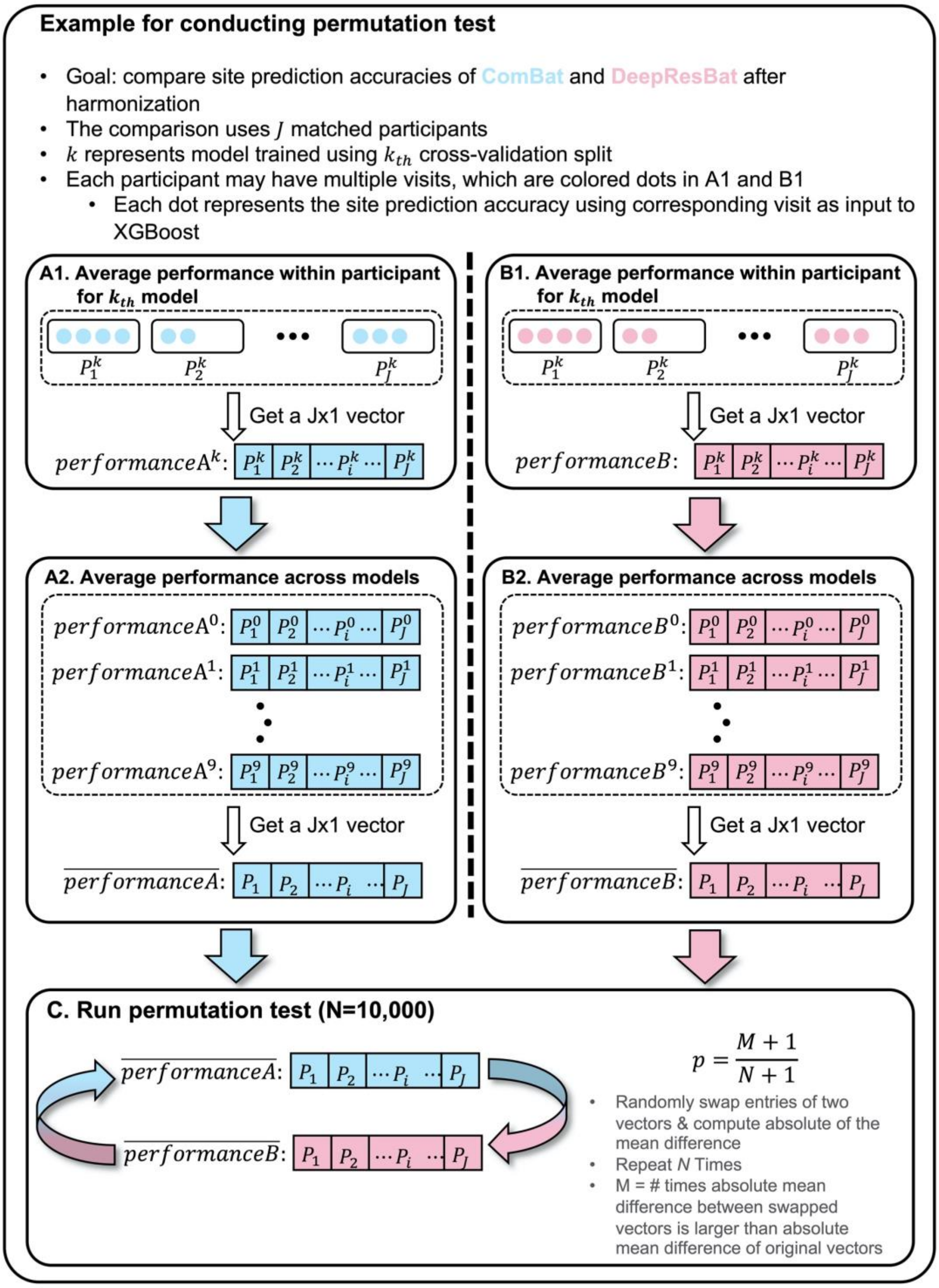


**Figure S4. Illustration of permutation test for comparing site prediction accuracies of ComBat and DeepResBat.** (A1) For a given model, we averaged the site prediction accuracies within each participant for ComBat. (B1) Same as A1 but for DeepResBat. (A2) Averaging the site prediction accuracies across the 10 models within each participant. (B1 & B2) Same as A1 and A2 but for DeepResBat. (C) Permute 10,000 times to obtain p value.

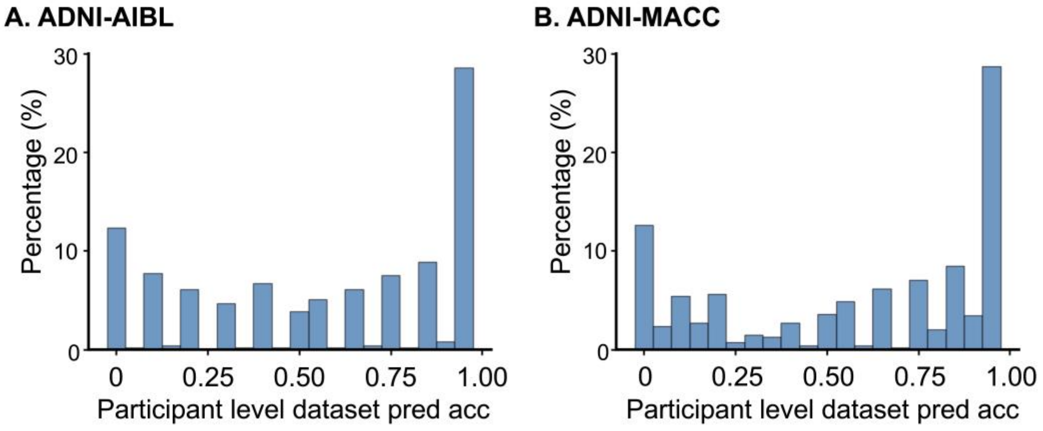


**Figure S5. Histogram of participant level dataset prediction accuracies on matched participants harmonized by DeepResBat**. (A) Histogram for ADNI-AIBL; (B) Histogram for ADNI-MACC.

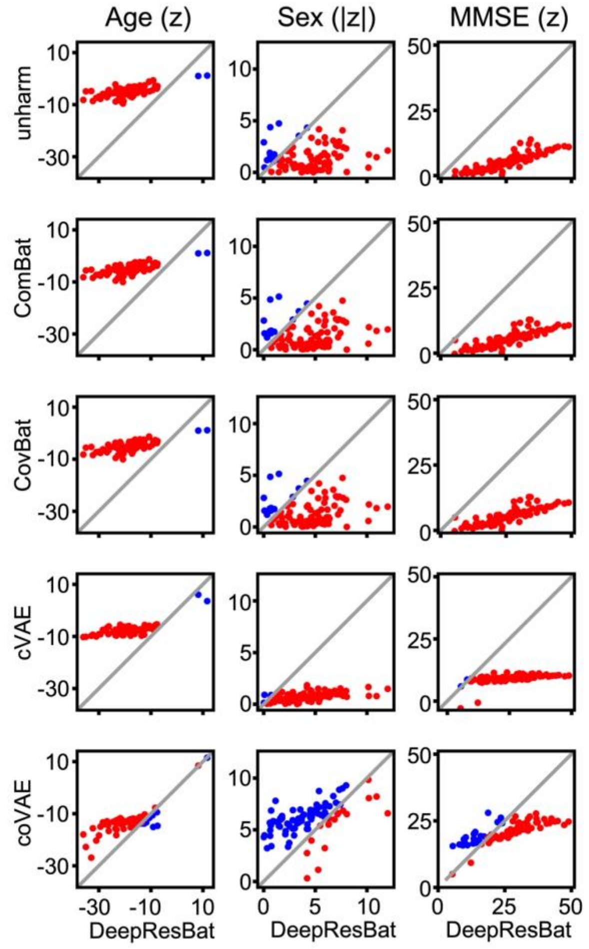


**Figure S6. Comparison of z statistics from GLM involving MMSE for DeepResBat and baselines on matched ADNI and AIBL participants.** Each row compares DeepResBat and one baseline approach: no harmonization (row 1), ComBat (row 2), CovBat (row 3), cVAE (row 4) and coVAE (row 5). Each column represents one covariate: age (column 1), sex (column 2), and MMSE (column 3). Each subplot compares z statistics of DeepResBat against another baseline for a given covariate across 87 grey matter ROIs. Each dot represents one grey matter ROI. Red dots indicate better performance by DeepResBat. Blue dots indicate worse performance by DeepResBat.

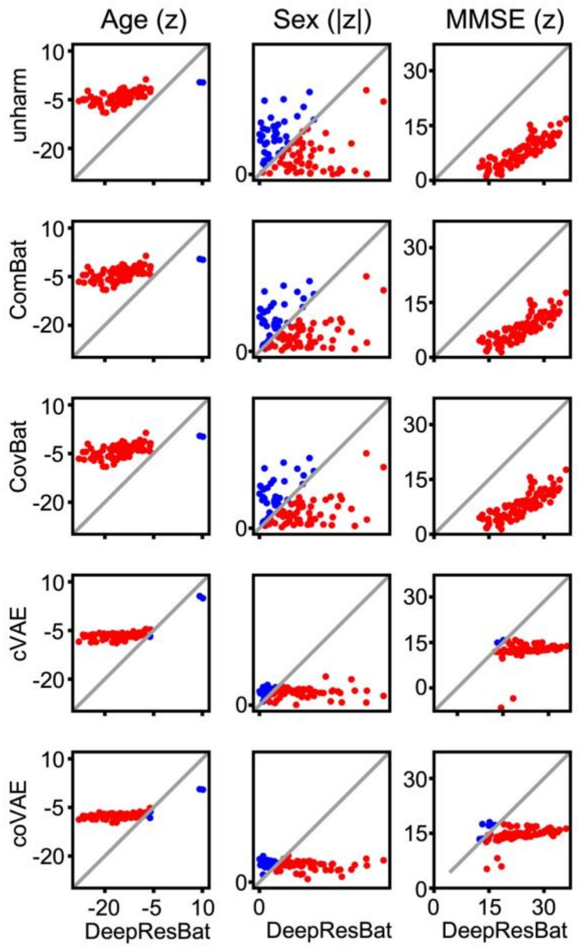


**Figure S7. Comparison of z statistics from GLM involving MMSE for DeepResBat and baselines on matched ADNI and MACC participants.** Each row compares DeepResBat and one baseline approach: no harmonization (row 1), ComBat (row 2), CovBat (row 3), cVAE (row 4) and coVAE (row 5). Each column represents one covariate: age (column 1), sex (column 2), and MMSE (column 3). Each subplot compares z statistics of DeepResBat against another baseline for a given covariate across 87 grey matter ROIs. Each dot represents one grey matter ROI. Red dots indicate better performance by DeepResBat. Blue dots indicate worse performance by DeepResBat.

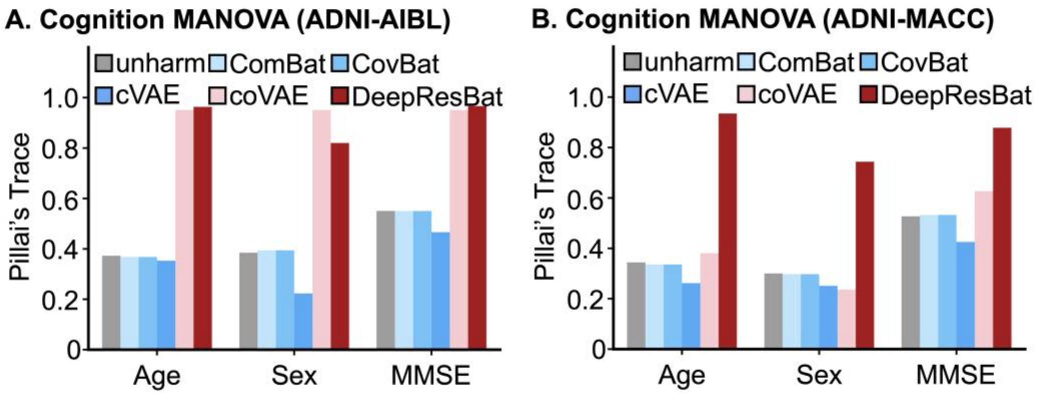


**Figure S8. Effect size of MANOVA involving MMSE.** A larger Pillai's Trace indicates a stronger association, and thus better performance. (A) Bar plot for matched ADNI and AIBL participants. (B) Bar plot for matched ADNI and MACC participants.

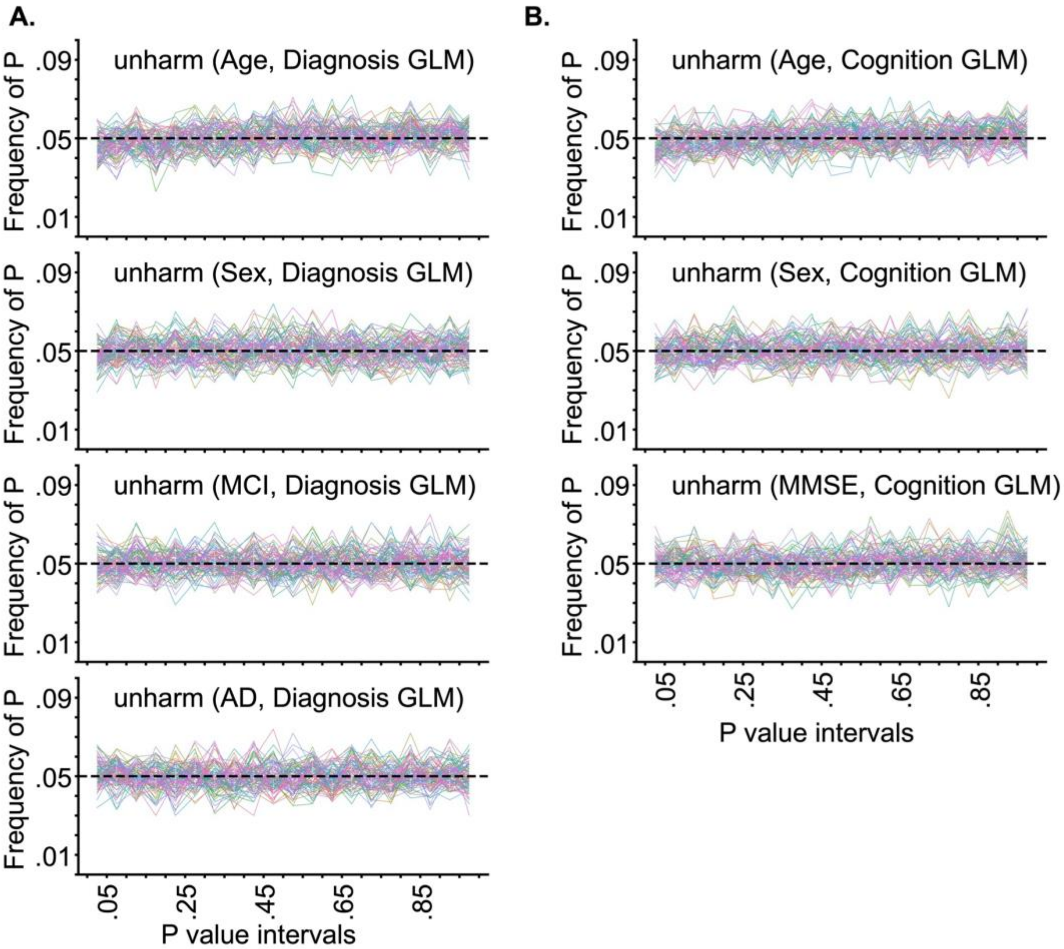


**Figure S9. Frequency of p values of unharmonized data for matched ADNI and AIBL participants by GLMs involving clinical diagnosis and MMSE based on 1000 permutations.** Each line corresponds to a single brain ROI. P values were binned in intervals of 0.05. Therefore, in the ideal scenario, the distributions of p values should follow a uniform distribution with a height of 0.05. (A) Frequency of p values by GLMs involving clinical diagnosis. (B) Frequency of p values by GLMs involving MMSE.

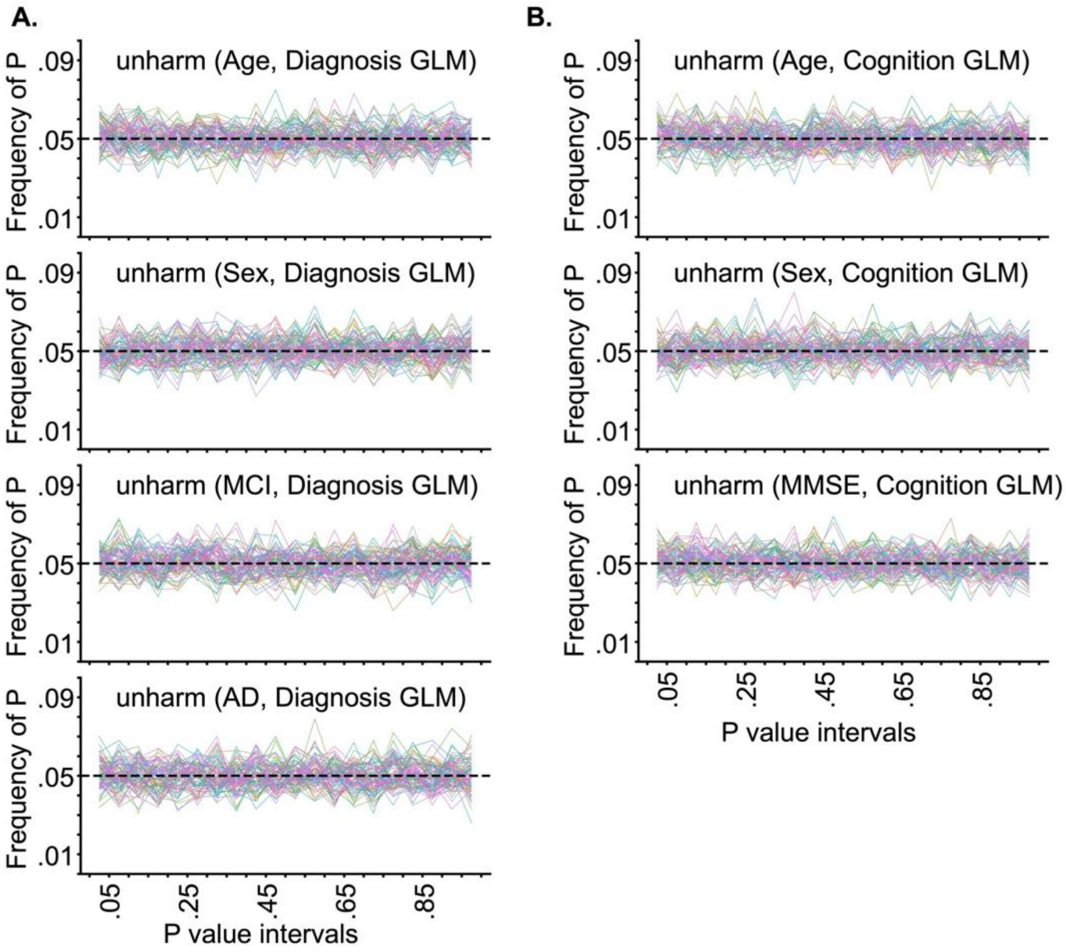


**Figure S10. Frequency of p values of unharmonized data for matched ADNI and MACC participants by GLMs involving clinical diagnosis and MMSE based on 1000 permutations.** Each line corresponds to a single brain ROI. P values were binned in intervals of 0.05. Therefore, in the ideal scenario, the distributions of p values should follow a uniform distribution with a height of 0.05. (A) Frequency of p values by GLMs involving clinical diagnosis. (B) Frequency of p values by GLMs involving MMSE.

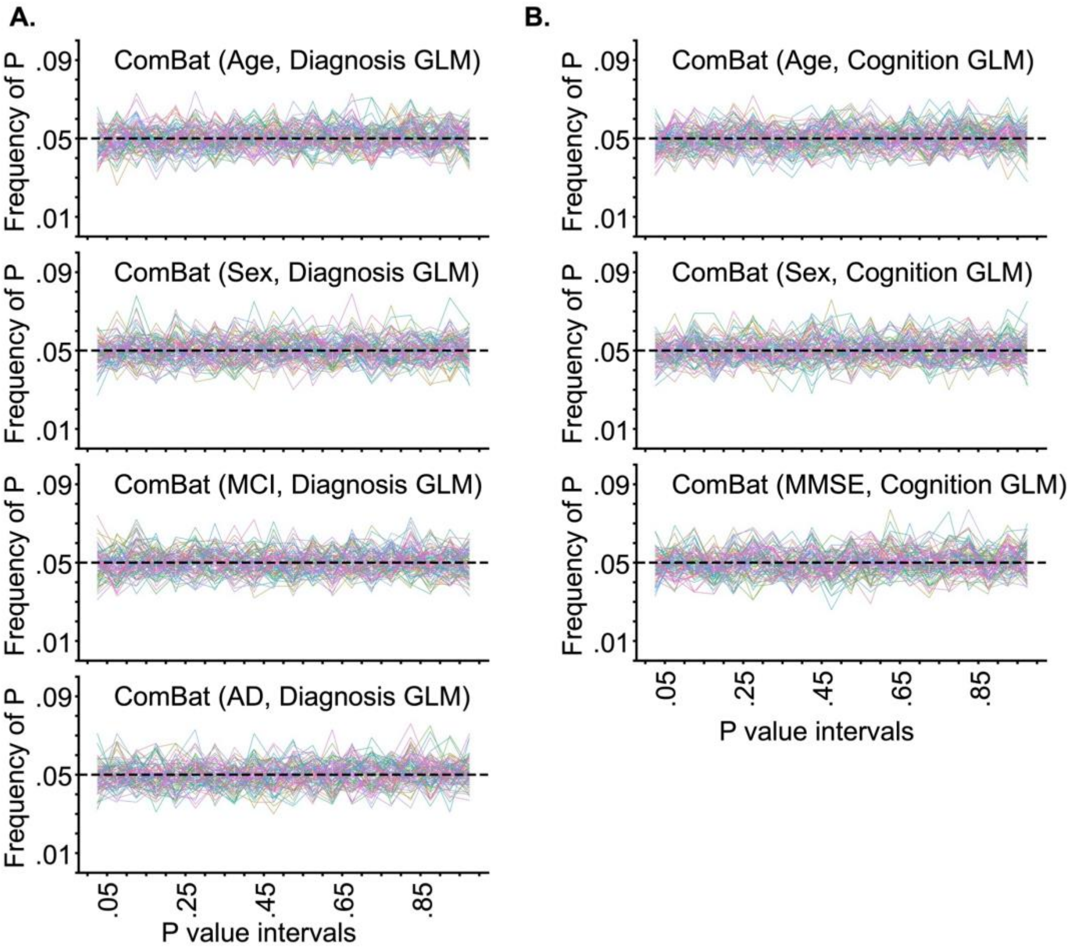


**Figure S11. Frequency of p values of ComBat for matched ADNI and AIBL participants by GLMs involving clinical diagnosis and MMSE based on 1000 permutations.** Each line corresponds to a single brain ROI. P values were binned in intervals of 0.05. Therefore, in the ideal scenario, the distributions of p values should follow a uniform distribution with a height of 0.05. (A) Frequency of p values by GLMs involving clinical diagnosis. (B) Frequency of p values by GLMs involving MMSE.

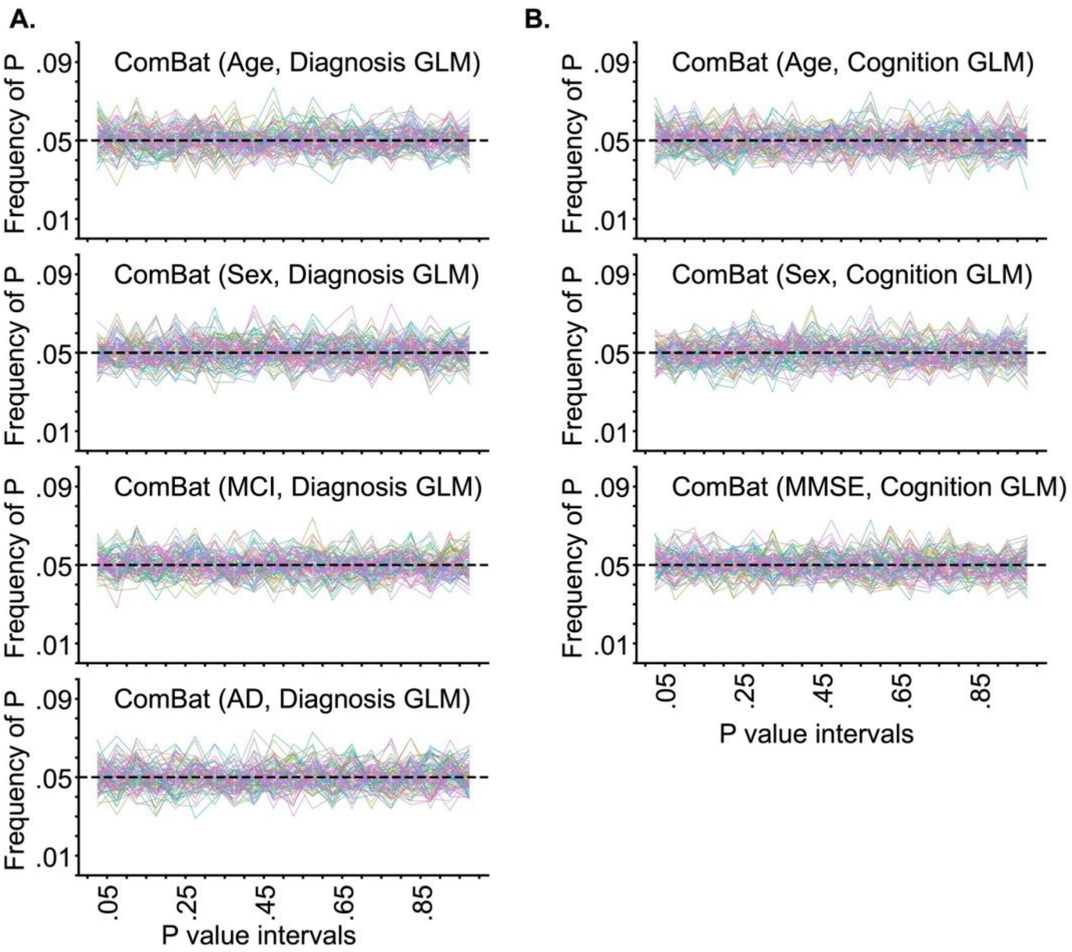


**Figure S12. Frequency of p values of ComBat for matched ADNI and MACC participants by GLMs involving clinical diagnosis and MMSE based on 1000 permutations.** Each line corresponds to a single brain ROI. P values were binned in intervals of 0.05. Therefore, in the ideal scenario, the distributions of p values should follow a uniform distribution with a height of 0.05. (A) Frequency of p values by GLMs involving clinical diagnosis. (B) Frequency of p values by GLMs involving MMSE.

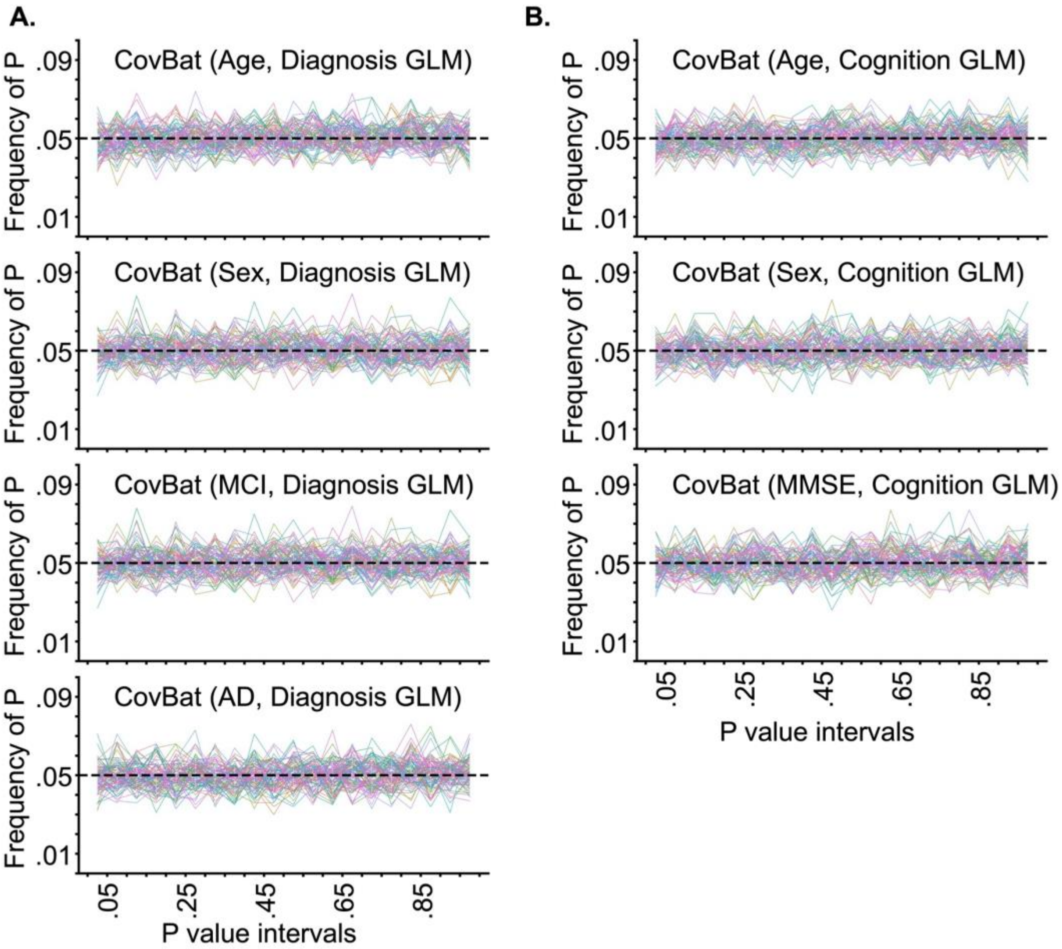


**Figure S13. Frequency of p values of CovBat for matched ADNI and AIBL participants by GLMs involving clinical diagnosis and MMSE based on 1000 permutations.** Each line corresponds to a single brain ROI. P values were binned in intervals of 0.05. Therefore, in the ideal scenario, the distributions of p values should follow a uniform distribution with a height of 0.05. (A) Frequency of p values by GLMs involving clinical diagnosis. (B) Frequency of p values by GLMs involving MMSE.

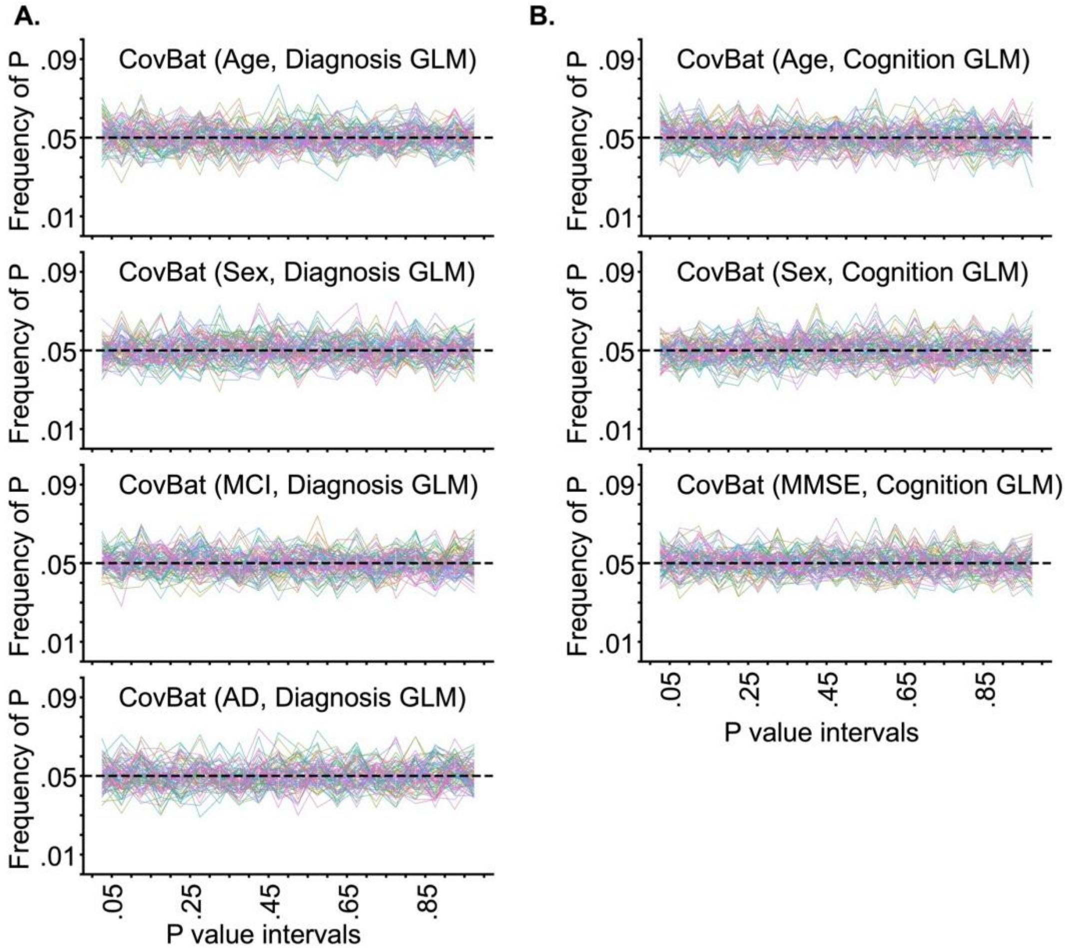


**Figure S14. Frequency of p values of CovBat for matched ADNI and MACC participants by GLMs involving clinical diagnosis and MMSE based on 1000 permutations.** Each line corresponds to a single brain ROI. P values were binned in intervals of 0.05. Therefore, in the ideal scenario, the distributions of p values should follow a uniform distribution with a height of 0.05. (A) Frequency of p values by GLMs involving clinical diagnosis. (B) Frequency of p values by GLMs involving MMSE.

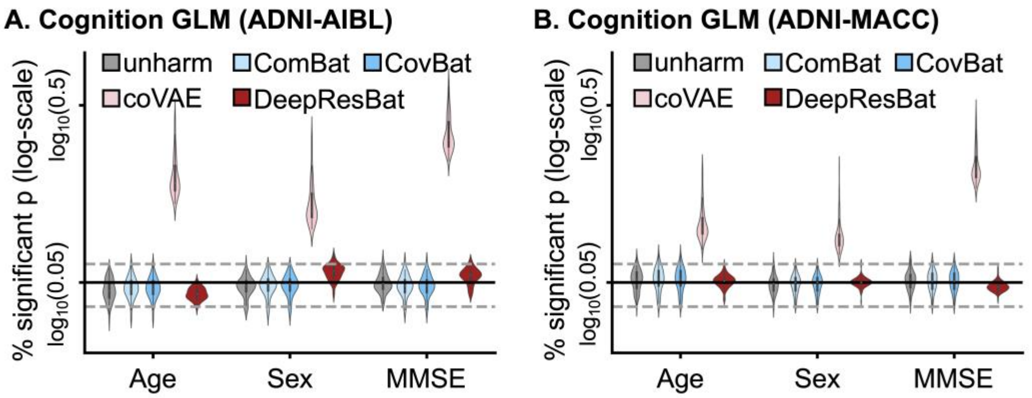


**Figure S15. Percentage of significant p values (i.e., $p < 0.05$) from GLM with MMSE after 1000 permutations of covariates.** More specifically, each data point in the violin plot represents a brain ROI volume. Percentage is calculated based on the number of permutations in which p value of corresponding covariate was significant (i.e., $p < 0.05$) divided by 1000 permutations. Percentage (vertical axis) is shown on a log scale. The black solid line is the expected percentage (which is 0.05), while the grey dashed lines indicated 95% confidence intervals. (A) GLM analysis involving MMSE for matched ADNI and AIBL participants. (B) GLM analysis involving MMSE for matched ADNI and MACC participants.

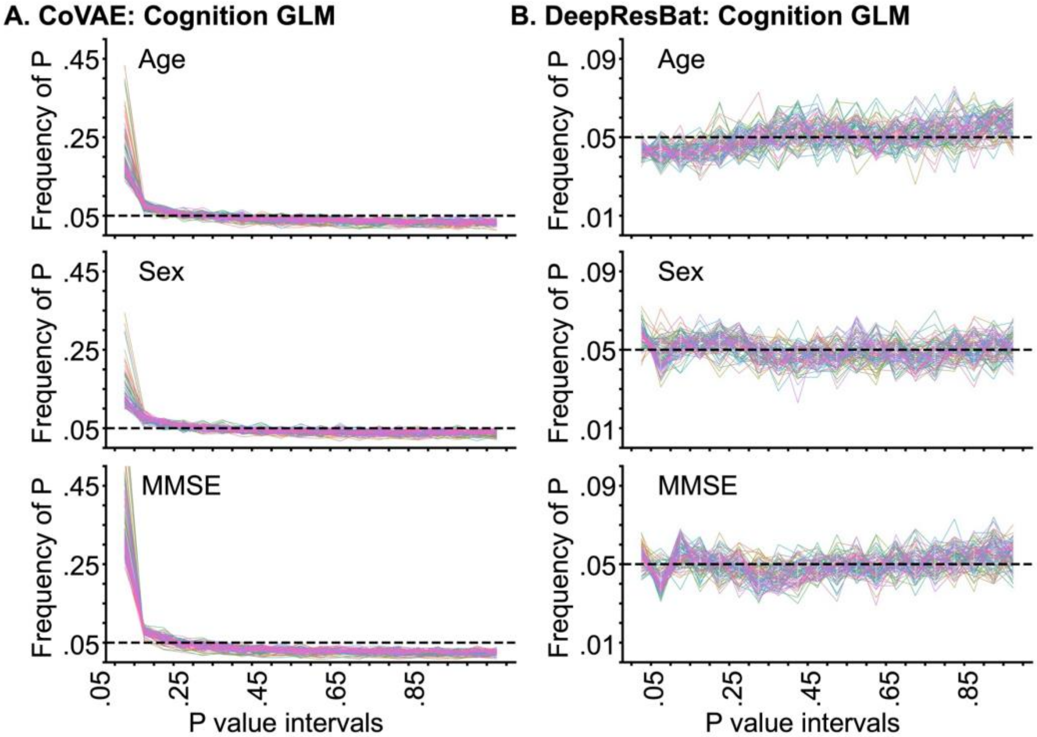


**Figure S16. Frequency of p values of coVAE and DeepResBat for matched ADNI and AIBL participants by GLM involving MMSE based on 1000 permutations.** Each line corresponds to a single brain ROI. P values were binned in intervals of 0.05. Therefore, in the ideal scenario, the distributions of p values should follow a uniform distribution with a height of 0.05. (A) Frequency of p values for coVAE. (B) Frequency of p values for DeepResBat.

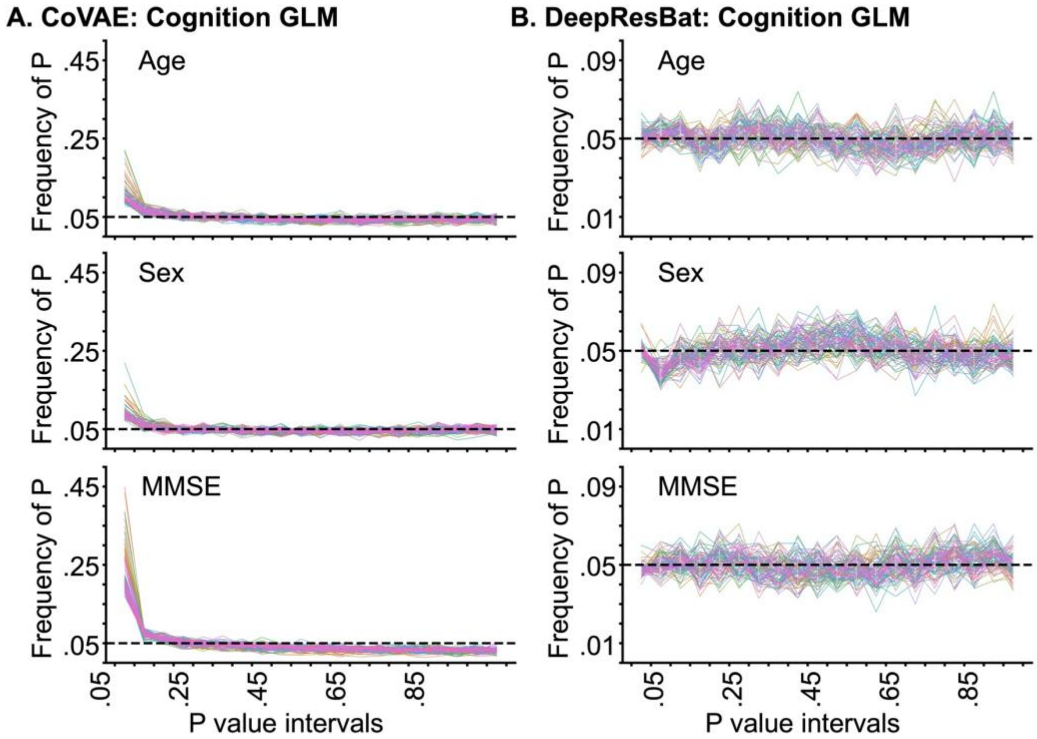


**Figure S17. Frequency of p values of coVAE and DeepResBat for matched ADNI and MACC participants by GLM involving MMSE based on 1000 permutations.** Each line corresponds to a single brain ROI. P values were binned in intervals of 0.05. Therefore, in the ideal scenario, the distributions of p values should follow a uniform distribution with a height of 0.05. (A) Frequency of p values for coVAE. (B) Frequency of p values for DeepResBat.

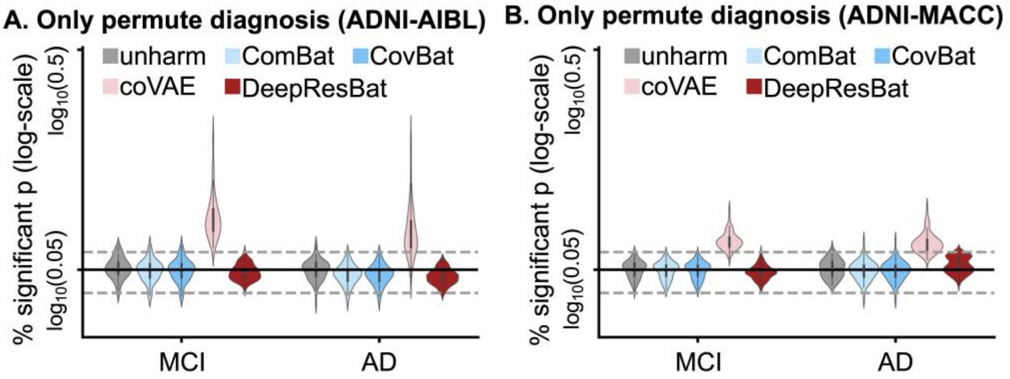


**Figure S18. Percentage of significant p values (i.e., $p < 0.05$) from GLM with clinical diagnosis after 1000 permutations of clinical diagnosis only.** More specifically, each data point in the violin plot represents a brain ROI volume. Percentage is calculated based on the number of permutations in which p value of corresponding covariate was significant (i.e., $p < 0.05$) divided by 1000 permutations. Percentage (vertical axis) is shown on a log scale. The black solid line is the expected percentage (which is 0.05), while the grey dashed lines indicated 95% confidence intervals. (A) GLM analysis involving clinical diagnosis for matched ADNI and AIBL participants. (B) GLM analysis involving clinical diagnosis for matched ADNI and MACC participants.

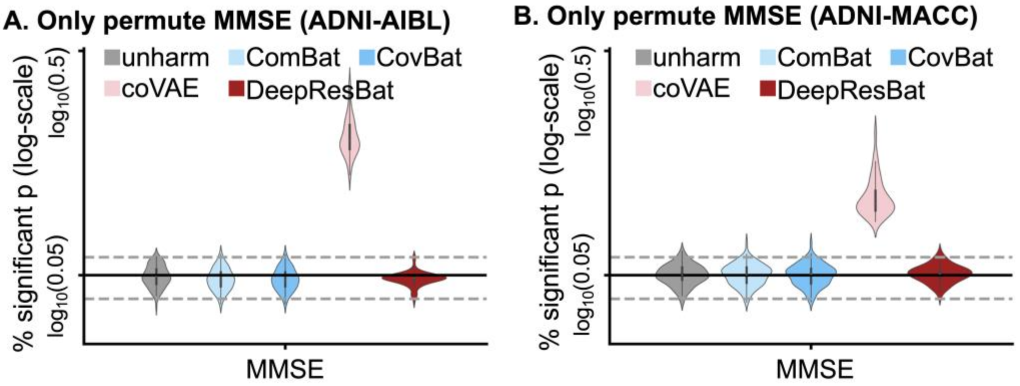


**Figure S19. Percentage of significant p values (i.e., $p < 0.05$) from GLM with MMSE after 1000 permutations of MMSE only.** More specifically, each data point in the violin plot represents a brain ROI volume. Percentage is calculated based on the number of permutations in which p value of corresponding covariate was significant (i.e., $p < 0.05$) divided by 1000 permutations. Percentage (vertical axis) is shown on a log scale. The black solid line is the expected percentage (which is 0.05), while the grey dashed lines indicated 95% confidence intervals. (A) GLM analysis involving MMSE for matched ADNI and AIBL participants. (B) GLM analysis involving MMSE for matched ADNI and MACC participants.

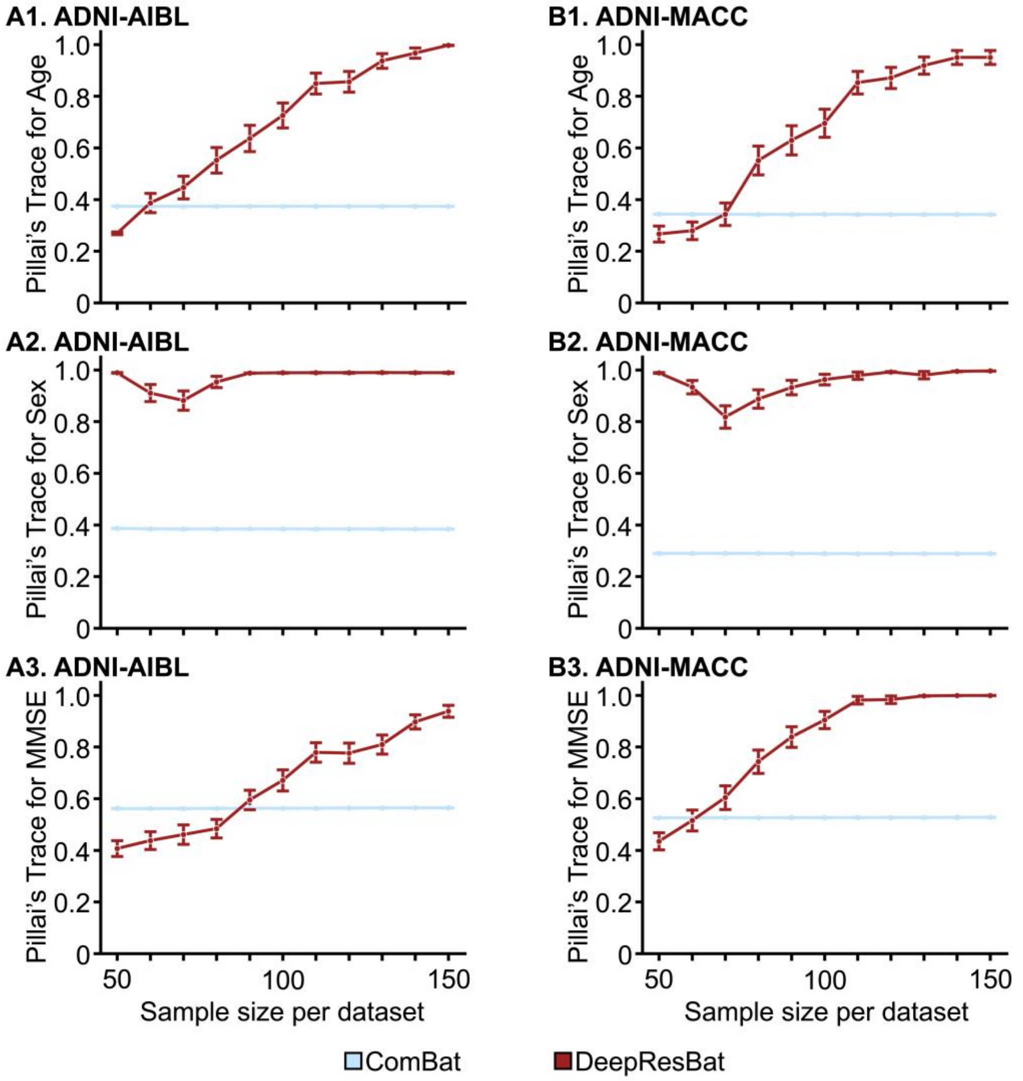


**Figure S20. Effect size bar plot by MANOVA involving MMSE for ComBat and DeepResBat with different sample sizes per site.** Each sampling was repeated 50 times, error bar is the standard error. Left column is result for sampling on ADNI-AIBL, right column is result for sampling on ADNI-MACC. Each row is effect size of interested variables. (A1) Effect size for association with age by MANOVA for ADNI-AIBL. (A2) Effect size for association with sex by MANOVA for ADNI-AIBL. (A3) Effect size for association with MMSE by MANOVA for ADNI-AIBL. (B1) to B(3): Same with (A1) to A(3) but for ADNI-MACC.

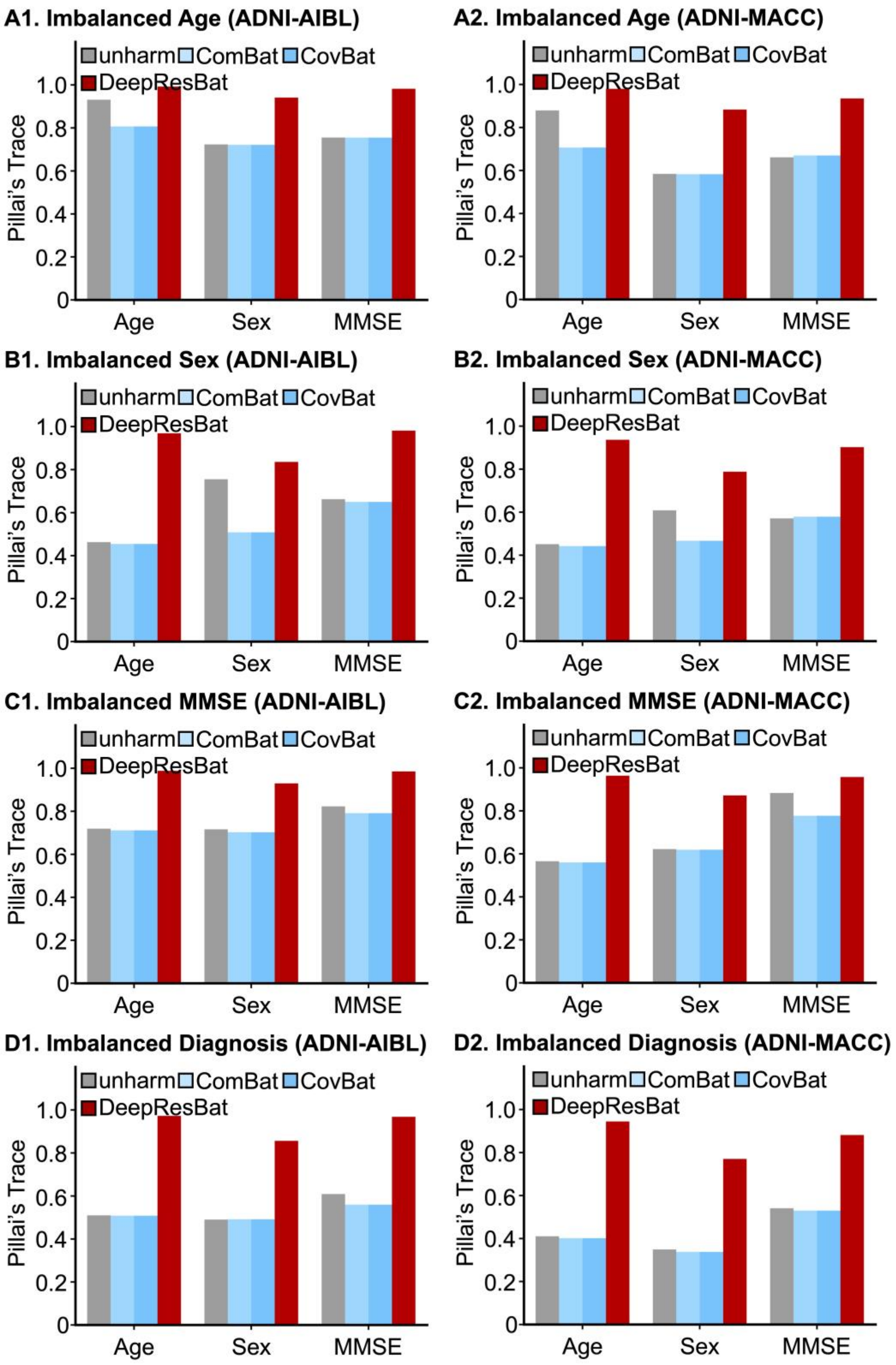


**Figure S21.** Effect size of MANOVA involving MMSE in test sets involving highly imbalanced covariate distributions. A larger Pillai's Trace indicates a stronger association, and thus better performance. The left column corresponding to harmonizing ADNI and AIBL. The right column corresponding to harmonizing ADNI and MACC. (A1) MANOVA effect sizes for ADNI-AIBL test set with highly imbalanced age distributions. (A2) Same as A1 but for ADNI and MACC. (B1) MANOVA effect sizes for ADNI-AIBL test set with highly imbalanced sex distributions. (B2) Same as B1 but for ADNI and MACC. (C1) MANOVA effect sizes for ADNI-AIBL test set with highly imbalanced MMSE distributions. (C2) Same as C1 but for ADNI and MACC. (D1) MANOVA effect sizes for ADNI-AIBL test set with highly imbalanced diagnosis distributions. (D2) Same as B1 but for ADNI and MACC.
